# Towards transboundary networks of climate-smart marine reserves in the Southern California Bight

**DOI:** 10.1101/2022.01.04.475006

**Authors:** Nur Arafeh-Dalmau, Adrian Munguia-Vega, Fiorenza Micheli, Ainoa Vilalta-Navas, Juan Carlos Villasenor-Derbez, Magdalena Précoma-de la Mora, David S. Schoeman, Alfonso Medellín-Ortíz, Kyle C. Cavanaugh, Oscar Sosa-Nishizaki, Theresa L.U. Burnham, Christopher J. Knight, C. Brock Woodson, Marina Abas, Alicia Abadía-Cardoso, Octavio Aburto-Oropeza, Michael W. Esgro, Noemi Espinosa-Andrade, Rodrigo Beas-Luna, Nirari Cardenas, Mark H. Carr, Katherine E. Dale, Frida Cisneros-Soberanis, Ana Laura Flores-Morales, Stuart Fulton, Emiliano García-Rodríguez, Alfredo Giron-Nava, Mary G. Gleason, Alison L. Green, Arturo Hernández-Velasco, Beatriz Ibarra-Macías, Andrew F. Johnson, Julio Lorda, Luis Malpica-Cruz, Gabriela Montaño-Moctezuma, Carolina Olguín-Jacobson, Alejandro Parés-Sierra, Peter T. Raimondi, Georgina Ramírez-Ortiz, Arturo Ramirez-Valdez, Héctor Reyes-Bonilla, Emily Saarman, Luz Erandi Saldaña-Ruiz, Alexandra Smith, Cecilia Soldatini, Alvin Suárez, Guillermo Torres-Moye, Mariana Walther, Elizabeth Burke Watson, Sara Worden, Hugh P. Possingham

**Affiliations:** Centre for Biodiversity and Conservation Science, School of Biological Sciences, The University of Queensland, St Lucia, Queensland, Australia; School of Earth and Environmental Sciences, The University of Queensland, St Lucia, Queensland, Australia; Desert Laboratory on Tumamoc Hill, University of Arizona, Tucson, AZ, 85721, USA; Applied Genomics Lab, La Paz, Baja California Sur, Mexico; Hopkins Marine Station and Stanford Center for Ocean Solutions, Pacific Grove, CA 93950, USA; Instituto de Investigaciones Oceanológicas, Universidad Autónoma de Baja California, Ensenada, Mexico; Bren School of Environmental Science & Management, University of California, Santa Barbara, Santa Barbara, CA, USA; Comunidad y Biodiversidad, A.C., Guaymas, Sonora., México; Global-Change Ecology Research Group, School of Science, Technology and Engineering, University of the Sunshine Coast, Maroochydore, QLD, Australia; Department of Zoology, Centre for African Conservation Ecology, Nelson Mandela University, Gqeberha, South Africa; Universidad Autónoma de Baja California, Facultad de Ciencias Marinas, Ensenada, Mexico; Department of Geography, University of California Los Angeles, Los Angeles, CA, USA; Centro de Investigación Científica y de Educación Superior de Ensenada (CICESE), Ensenada, Baja California, Mexico; Department of Wildlife, Fish, and Conservation Biology, University of California, Davis, Davis, CA 95616, USA; Coastal and Marine Institute and Department of Biology, San Diego State University, San Diego, CA 92106, USA; College of Engineering, University of Georgia, Athens, GA, USA; Universidad Autónoma de Baja California Sur, Carretera al Sur 5.5., La Paz C.P 23080, Mexico; Scripps Institution of Oceanography, University of California San Diego, La Jolla, CA, USA; California Ocean Protection Council, Sacramento, California, USA; The Nature Conservancy (Mexico), Mexico; Department of Ecology and Evolutionary Biology, University of California Santa Cruz, Santa Cruz, California, USA; Stanford Center for Ocean Solutions, Pacific Grove, CA 93950, USA; The Nature Conservancy, Sacramento, CA, United States; Red Sea Research Centre, King Abdullah University of Science and Technology, Saudi Arabia; MarFishEco Fisheries Consultants Ltd, Edinburgh, UK; The Marine SPACE Group, The Lyell Centre, Institute of Life and Earth Sciences, School of Energy, Geoscience, Infrastructure and Society, Heriot-Watt University, Edinburgh, UK; Australian Rivers Institute, School of Environment and Science, Griffith University, Southport, Queensland, Australia; Centro de Investigaciones Biológicas del Noroeste, Av. Instituto Politécnico Nacional 195, CP 23096, La Paz, Baja California Sur, Mexico.; Centro de Investigación Científica y de Educación Superior de Ensenada (CICESE), La Paz, Baja California Sur, Mexico.; Centro de Estudios Biológicos, Medio Ambiente y Recursos Naturales, A.C., Felipe Carrillo Puerto, Quintana Roo, México; Department of Biodiversity, Earth and Environmental Sciences, Drexel University, Philadelphia, PA 19104, USA; California Department of Fish and Wildlife, Marine Region, 350 Harbor Boulevard, Belmont, California, 94002 USA

## Abstract

Climate-smart conservation addresses the vulnerability of biodiversity to climate change impacts but may require transboundary considerations. Here, we adapt and refine 16 biophysical guidelines for climate-smart marine reserves for the transboundary California Bight ecoregion. We link several climate-adaptation strategies (e.g., maintaining connectivity, representing climate refugia, and forecasting effectiveness of protection) by focusing on kelp forests and associated species. We quantify transboundary larval connectivity along ∼800 km of coast and find that the number of connections and the average density of larvae dispersing through the network under future climate scenarios could decrease by ∼50%, highlighting the need to protect critical steppingstone nodes. We also find that although focal species will generally recover with 30% protection, marine heatwaves could hinder subsequent recovery in the following 50 years, suggesting that protecting climate refugia and expanding the coverage of marine reserves is a priority. Together, these findings provide a first comprehensive framework for integrating climate resilience for networks of marine reserves and highlight the need for a coordinated approach in the California Bight ecoregion.

## Introduction

Marine reserves can rebuild the biomass of overfished species^1^, conserve biodiversity^2^, and enhance the resilience and adaptive capacity of ecosystems to climate impacts^3–7^. However, delivering large-scale benefits requires networks of marine reserves that are functionally interconnected, and are large enough to protect the underlying biophysical processes that maintain species distribution and composition^8^. A rich literature exists on biophysical guidelines for designing networks of marine reserves for fisheries, conservation, and climate-adaptation objectives^9–12^, but most are limited to analysis within country or state boundaries. By contrast, ecoregion-scale planning efforts may span thousands of kilometres and, in many cases, cross multiple national or international jurisdictions^13, 14^. Consequently, before designing networks of transboundary marine reserves, planners need to develop shared biophysical guidelines and comprehensive spatial analyses across borders^15^.

Climate change is one of the main threats to marine ecosystems^16^ that could be partially addressed by large-scale, coordinated, climate-smart networks of marine reserves^11, 17, 18^. Although marine reserves cannot directly abate climate-change threats, they can indirectly mitigate climate-change impacts by promoting ecological resilience^3–5, 19^. Climate-smart conservation is a multiple-step approach that addresses the vulnerability of species and ecosystems to changes in climate and ocean chemistry, and supports resilience of populations and ecosystems^17^. Climate-adaptation strategies include protecting areas that act as climate refugia^11, 18, 20^, maintain ecological connectivity to ensure metapopulation persistence^21^, facilitate species’ range shifts, and recover important species for ecosystem functioning (e.g., predators)^22^. Complementary strategies include supporting ecosystem resilience by addressing threats not directly abated by marine reserves^11^ (e.g., pollution and other impacts) and protecting and restoring habitats that could mitigate some of the effects of climate change (e.g., through carbon sequestration). Because global conservation targets aim to protect 30% of marine habitats by 2030^23, 24^ while adapting to climate change, there is a need to integrate multiple adaptation strategies for designing networks of climate-smart reserves.

Given the highly dynamic nature of the oceans, biophysical modelling of larval dispersal is an essential tool to inform transboundary conservation^25, 26^. However, these models should consider the implications of climate change on larval dynamics, such as changes in dispersal distances and the availability of suitable habitats for settlement^21, 27^. These considerations are essential because transboundary dispersal may be critical for metapopulation persistence. Notably, certain areas may be less impacted by climate change and act as climate refugia^28^, providing food, shelter, and habitat, despite future changes. Identifying climate refugia at large spatial scales can be challenging, requiring the use of ecosystem attributes (resistance, resilience, persistence^29^) or environmental proxies (e.g., micro climates)^30^. If we map these areas, we can prioritize their protection and assess changes in larval connectivity for future scenarios. Climate change may undermine the effectiveness of transboundary networks of marine reserves to facilitate recovery of exploited species. Thus, it is a priority to assess whether proposed protection targets^23, 24^ will facilitate recovery of overexploited species in the future and whether increased protection or alternative strategies will be necessary to ensure their recovery.

The Southern California Bight ecoregion (henceforth “California Bight”) in the northwest Pacific Ocean ─ shared between the state of California, USA, and the Peninsula of Baja California, Mexico ─ has a long history of research cooperation and binational environmental agreements^31, 32^. It is considered a marine climate-change “hotspot” – rapidly warming ocean regions that are natural laboratories for evaluating climate adaptation options^33^. Recent marine heatwaves^34–37^ and prolonged hypoxic events^3^ exemplify the impacts of climate variability and environmental extremes on species, ecosystems, and coastal economies of this region. Documentation of these changes includes mass mortality events and range shifts of economically or ecologically important species^3, 34, 37–39^. This threatens the conservation and economic sustainability outcomes that both countries seek to deliver through marine zoning.

In 2012, California implemented a network of marine protected areas covering 16% of state waters, with more than half being fully protected marine reserves. This network was based on stakeholder input and biophysical guidelines to ensure ecological connectivity, habitat representation and replication, while balancing users’ needs^40^. However, the establishment of marine protected areas did not include climate-adaptation objectives^41^. Moreover, the network did not consider the transboundary nature of the region, where many species move across the USA-Mexico border through larval dispersal and adult movement.

By contrast, although Baja California has extensive coastal areas protected, there is a lack of an integrated network of marine reserves, with less than 1% of the coastal waters fully protected^20, 42^. In this region, coastal resource management is based on territorial user rights granted to fishing cooperatives and independent permit holders, with successful local cases of community-based marine reserves sparking other cooperatives’ interest^42^. Additional community-led marine reserves are awaiting government approval, while environmental NGOs, scientists, fishing cooperatives, and governmental agencies are promoting a region-wide marine spatial planning process^42^.

Here, we first adapt and refine 16 biophysical guidelines for climate-smart, transboundary marine reserve design for the California Bight ecoregion ─ spanning over eight degrees of latitude. We used kelp forest ecosystems (dominated by giant kelp, *Macrocystis pyrifera*) and focal species of fishes and invertebrates of commercial and ecological importance to identify, analyze, and map areas that integrate and meet the proposed climate-smart guidelines. Then we explore the magnitude and importance of transboundary connectivity in the region and ask whether binational connections will be lost under future climate scenarios. We also ask whether sea surface temperature variability can be used as a proxy for climate refugia and whether marine heatwaves in the following 50 years will undermine the effectiveness of marine reserves for facilitating recovery of vulnerable species in the California Bight. Our research can inform the delivery of networks of climate-smart marine reserves by 2030 in the California Bight and other regions.

## Methods

### Study area

The California Bight ecoregion is located in the southern California Current System in the northeast Pacific Ocean and spans the USA-Mexico international border, from Point Conception, California, USA, in the north to Punta Abreojos, Baja California Sur, Mexico, in the south^43^. This highly productive ecoregion is in a transitional zone between the southward-flowing, cold, nutrient-rich California Current and the northward-flowing, warm, nutrient-depleted Davidson Current^44^. This transboundary ecoregion is characterized by strong latitudinal gradients in environmental conditions and oceanographic features that support a diverse assemblage of species and habitats^45^. We divided the California Bight into four sub-regions: southern California, northern and central Baja California, and Guadalupe Island. These four sub-regions represent geographic borders (the USA-Mexico border) and distinct biogeographic areas where species composition varies because of environmental conditions^46^.

### Developing and integrating biophysical guidelines

We developed biophysical guidelines (Table 1) for designing transboundary networks of climate-smart marine reserves for biodiversity conservation, fisheries replenishment, and climate-adaptation objectives in the California Bight. We compiled, adapted, and refined the guidelines using criteria developed for California, Mexico, and other regions^9, 10, 40, 47^. Our work does not provide an extensive review of the proposed guidelines or their ecological rationale, as this has been addressed in previous work^9, 10^. Instead, we summarise our findings in the context of transboundary and climate-smart spatial planning.

**Table 1.**
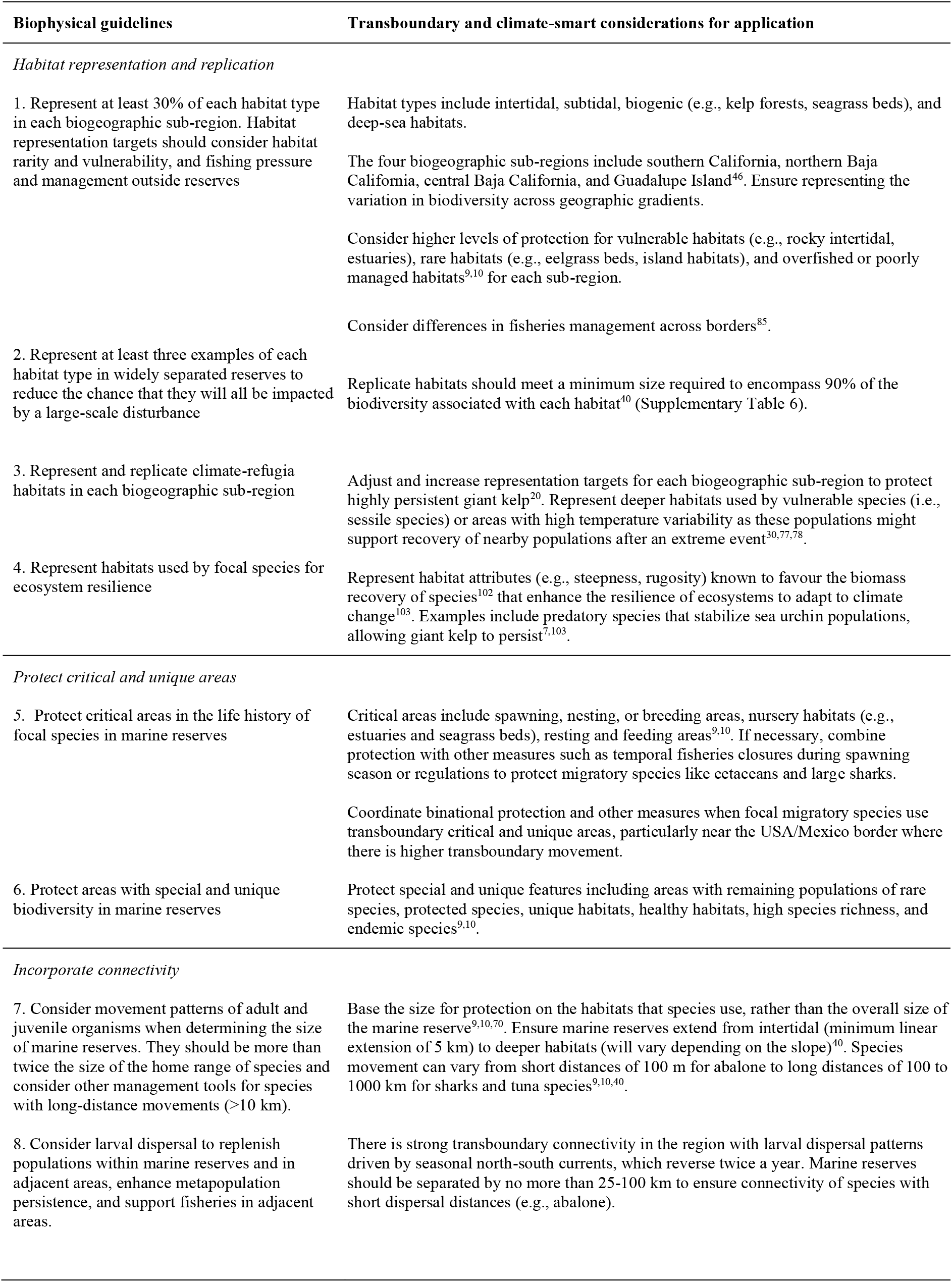
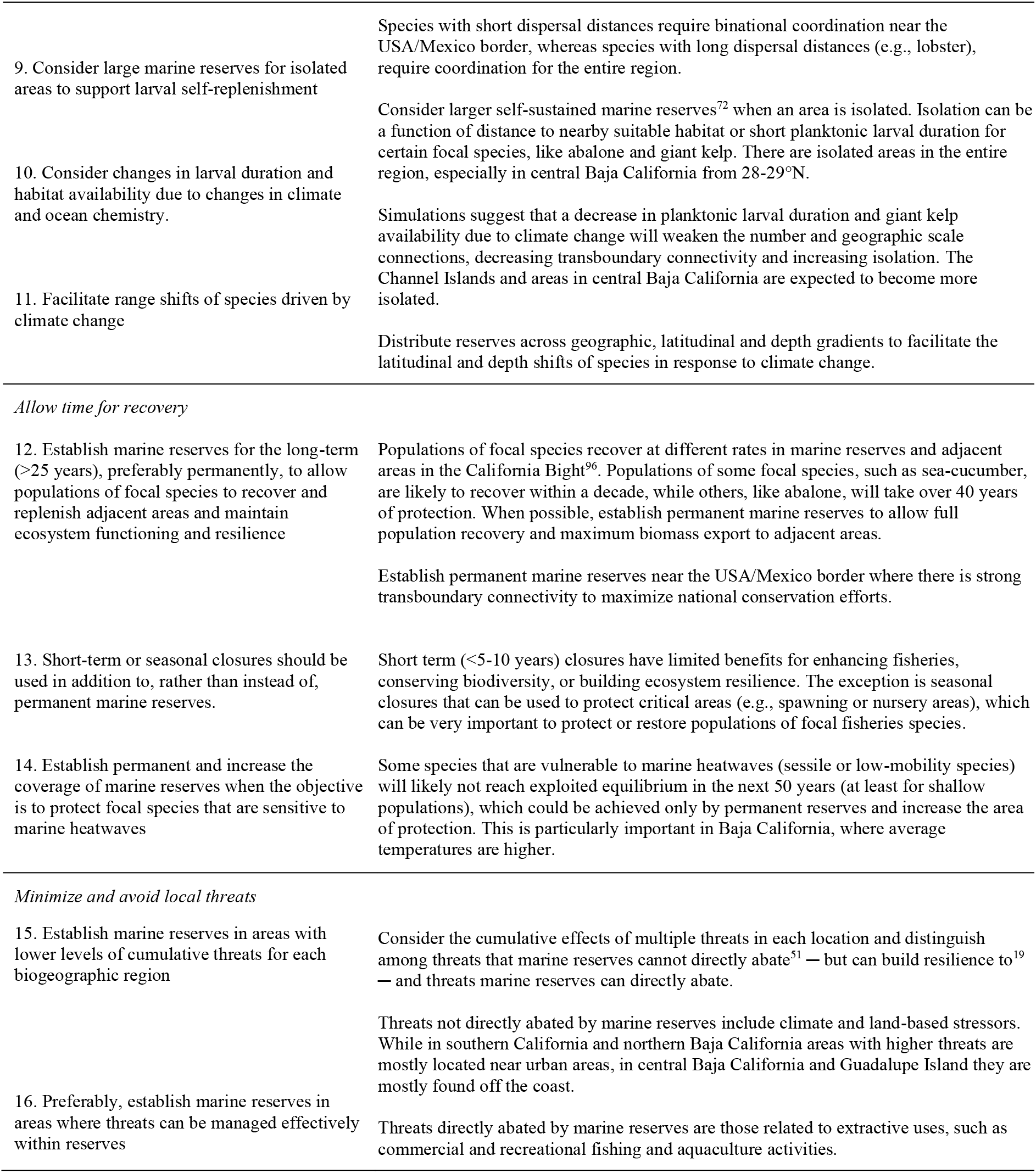
Biophysical guidelines and transboundary and climate-smart considerations for the design of networks of marine reserves in the Southern California Bight, compiled, adapted, and refined from California, Mexico, and other regions^9, 10, 40^.

Our work builds on a workshop held in 2017, which developed biophysical design guidelines for the Pacific region of Baja California^42^. Following this first workshop, five working groups comprising researchers from Mexico, the USA and Australia, fishing cooperatives, governmental agencies, and non-profit organizations in Mexico and the USA further developed the principles and their application to the California Bight (Table 1). Finally, following a second workshop held in 2019, we conducted spatial analysis and developed maps that integrate and meet some of the proposed guidelines for the California Bight, focussing on giant kelp (*M. pyrifera*) forests (henceforth “giant kelp”) and six focal species of commercial and ecological importance associated with giant kelp. Focal species included California Sheephead (*Semicossyphus pulcher*), sea basses (*Paralabrax clathratus* and *P. nebulifer*), whitefish (*Caulolatilus princeps*), spiny lobster (*Panulirus interruptus*), abalone (*Haliotis* spp.), sea urchins (*Mesocentrotus franciscanus* and *Strongylocentrotus purpuratus*), and sea cucumbers (*Apostichopus* spp. And *Parastichopus* spp.). We did not always use the same focal species among principles and analysis (see, Supplementary Table 1) because of differences in the availability of data, and in some cases, we used data from multiple taxa as composite focal species.

The proposed transboundary climate-smart guidelines fall into five major categories: (1) habitat representation and replication; (2) protecting critical and unique areas; (3) incorporating connectivity; (4) allowing time for recovery; and (5) minimizing and avoiding threats (Fig. 1). Marine habitats are broad indicators of the distribution of biodiversity^48^, which are relatively easy to map, and if adequately represented (e.g., protect enough of each habitat) can effectively protect biodiversity and replenish overfished populations^9, 10^. Moreover, each habitat should be replicated in the network in case a large-scale disturbance impacts part of the system^49^. It is also important to represent and replicate habitats or areas more resistant and resilient to climate stressors (climate refugia), which can replenish impacted populations and habitats^9, 10^.

**Fig. 1.**
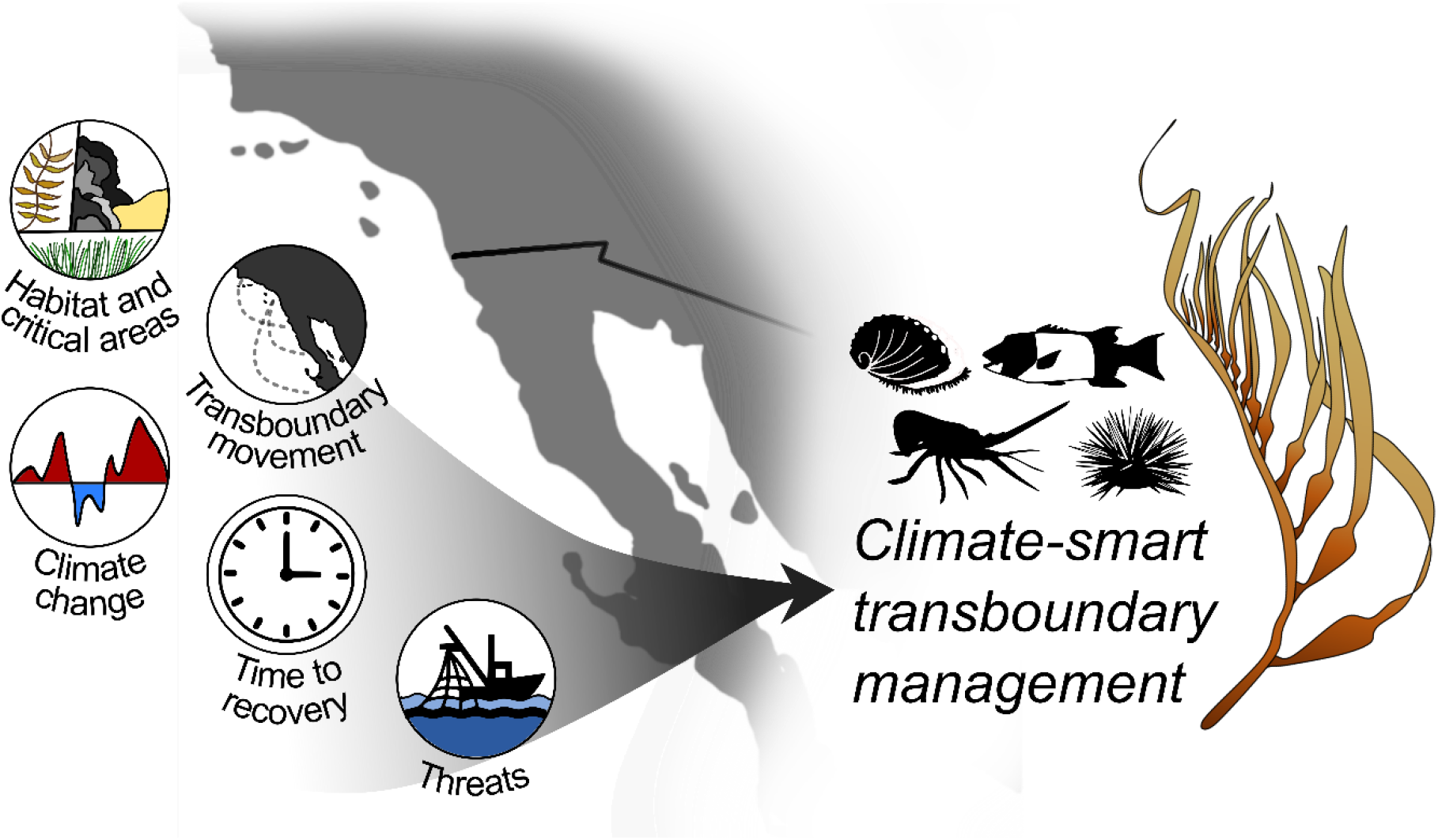
Graphical framework. Represents the biophysical principles used to develop spatial maps and analyses based on giant kelp forest ecosystems and associated species for transboundary networks of climate-smart marine reserves in the California Bight.

To facilitate recovery of focal species populations, the network should also protect areas critical for their life cycles, such as those required for spawning, nesting, or breeding^9, 10^. Notably, because most marine species are structured as metapopulations, connected through movements of adults, juveniles, or larval dispersal, ensuring the metapopulation persistence of focal species requires incorporating ecological connectivity^9, 10, 47, 50^. It is also crucial to consider changes in ocean conditions that could modify patterns of larval dispersal and recruitment and precipitate range shifts among species. Determining how much time overexploited species need to recover is essential to assessing the network’s effectiveness, especially if this is done while evaluating the projected impacts of climate change for vulnerable species. Finally, we need to consider threats that marine reserves can abate, versus those that reserves cannot abate, therefore requiring alternative strategies to support ecosystem resilience^19, 51^.

### Binational habitat mapping

We mapped the distribution of intertidal, estuarine, subtidal, and deep-sea habitats in the territorial seas (within 12 nautical miles of the coast) of the California Bight (Supplementary Table 2 and 3). We extracted depth contours (30, 100, and 200 m) from the General Bathymetric Chart of the Oceans^52^ and the California Department of Fish and Wildlife (CDFW, 2010 version) using ESRI ArcGIS Pro v10.8. We then used these contours to classify subtidal habitats based on depth categories (0-30, 30-100, 100-200, over 200 m). We obtained intertidal, estuarine and subtidal habitat polygons for southern California (CDFW, 2010 version) and northern Baja California from published work^53^. There was no available coastline or habitat mapping in the sub-regions of central Baja California and Guadalupe Island. We followed Arafeh-Dalmau, et al. ^53^ and digitized the coastline and mapped intertidal and subtidal habitats by visualizing Google Earth historical images.

We then combined existing maps of giant kelp distribution for California (CDFW, 2010 version) and Baja California^53^ with a 35-year satellite time series that maps the distribution and persistence of giant kelp at 30-m^2^ grid resolution, and classifies giant kelp persistence into three classes (high, mid, and low). For more details see Arafeh-Dalmau, et al. ^20^. Highly persistent giant kelp can be a good proxy for climate refugia because they have endured through time despite multiple cycles of marine climate oscillations^20^. We classified giant kelp polygons that did not overlap with persistence maps as having low persistence. We also collected information and mapped the distribution of tidal flats in southern California^54^ and eelgrass (*Zostera marina*) for northern and central Baja California from existing information (Pronatura Noroeste A.C.) and *in situ* presence-absence surveys combined with low-altitude drone imagery (for more details, see Supplementary Table 3). Finally, we catalogued geomorphic features data^55^ to map submarine canyons.

### Vulnerability of habitats and focal species to human threats

We conducted an expert-knowledge survey to quantify the vulnerability of six marine habitat types (estuaries, rocky intertidal, seagrass, kelp forest/rocky reefs, deep sea and pelagic) to eight major threats (marine heatwaves, ocean acidification, hypoxia, sea-level rise, storms, resource extraction, pollution, and physical habitat destruction) for the California Bight. The survey allowed experts to score the vulnerability of each habitat to each threat. Options included no threat, low, moderate, and high threat (coded 0, 1, 2, and 3, respectively; for more details see Supplementary Methods).

We complemented and expanded the expert-knowledge survey through a review of published studies for the California Bight that empirically examined the vulnerability and recovery rate of five focal species (California Sheephead, lobster, abalone, sea urchin, giant kelp; Supplementary Table 1) following marine heatwaves, hypoxia, and ocean acidification. We scored vulnerability to climate stressors that a species experiences as high for lethal effects, medium for extensive sublethal effects, and low for limited sublethal effects. We scored a species recovery rate from climatic stresses as rapid (< 1 year), moderate (2-5 years), or slow (> 5 years) (for more details, see Supplementary Methods).

### Sea surface temperature variability as a proxy for climate refugia

To identify potential climate refugia, we used two proxies: temperature variability^30^ and giant kelp persistence^20^. We analyzed patterns in their spatial variation and their correlation across a 1-km^2^ grid in the California Bight. We conducted this analysis to ascertain whether sea surface temperature variability is a comprehensive proxy of climate refugia for each sub-region in the California Bight. We obtained 17 years (2003-2019) of daily SST data from the Aqua-MODIS satellite at 1-km^2^ grid resolution, accessed through the NOAA-ERDDAP data repository (https://coastwatch.pfeg.noaa.gov/erddap/index.html). We estimated an annual cycle using a 30-day weighted moving average smoothing window for each year to create an annual climatology. We then used these annual cycles to estimate SST variability by computing the variance relative to the moving mean of the annual cycle (see Supplementary Fig. 1, for example). This SST variability metric represents high-frequency (24-h or faster) temperature variability. We then computed the mean SST variance across the 1-km^2^ grid to evaluate the accuracy of our model for SST variability with *in situ* sensors (a series of MiniDOT temperature/dissolved-oxygen sensors [PME Inc.] across Baja California^56^) and compared variability at sub-daily intervals. We then computed the mean giant kelp persistence^20^ over the same 1-km^2^ grid cells. We examined the correlation (linear least-squares regression) between these datasets for each sub-region, excluding Guadalupe Island, which does not have giant kelp.

### Transboundary movement of juveniles and adults into critical areas

To assess transboundary connectivity, we reviewed and mapped the transboundary movements of sharks, bony fishes, seabirds, marine mammals, and sea turtles between southern California and the three sub-regions in Baja California. We reviewed data from published papers and public databases (e.g., Tagging of Pacific Predators (TOPP) animal tracking dataset) for the period 1999-2019 using a combination of keywords: (1) movement or migration, (2) adults or juvenile, and (3) shark, bony fish, sea bird, marine mammals, or sea turtles. Because most studies tagged animals in USA waters, we considered connectivity only for organisms released in USA waters that moved into Mexican juvenile or adult habitats (e.g., spawning, breeding, nursery, resting, or foraging habitats). We aggregated the number of tagged studies that released organisms in southern California that were subsequently recorded in the three sub-regions in Baja California into 0.25° grid cells. Grid cells with high numbers of tagged animals recorded in Baja California were identified as essential areas for juvenile and adult transboundary connectivity.

### Transboundary larval dispersal model and impact of climate change

To further explore transboundary connectivity, we implemented the AGRIF version of the Regional Ocean Modelling System (ROMS; for more details, see Supplementary Methods, Supplementary Fig. 2) to simulate passive spores and larval dispersal (henceforth “larval dispersal”) for four focal species (California Sheephead, abalone, sea urchin and giant kelp; Supplementary Table 1) in the California Bight. We obtained information about spawning time and planktonic larval duration (PLD) from the literature (Supplementary Table 4), and for simplicity rounded to the nearest week when PLD exceeded 7 days.

To measure larval connectivity, we divided the coast into 54 polygons, each covering 20 km of latitude and limited by the 200 m isobath (Supplementary Fig. 3). This isobath represents the edge of the continental shelf that is a limit for coastal environments, where most fishing takes place^43^. In the centroid of each polygon, we released 1,000 virtual larvae at the start of each month of the year and followed their trajectories hourly for 60 days (2 months). We imported hourly coordinates for each modelled particle into MATLAB (Mathworks). We identified the intersection between particles and each polygon at the end of the PLD with a selection-by-location function. We generated connectivity matrices reflecting the proportion of larvae that settled in each polygon relative to the total number of larvae released at each site. We averaged matrices for the larval release dates within each month during each species’ spawning season. We calculated local retention as the proportion of larvae released within a polygon remaining within the natal area at the end of the PLD for each species. We explored connectivity matrices for each season using graph theory and a spatial network approach using the software GEPHI^57^, where nodes represent larval release sites and links represent directional larval dispersal probabilities. We estimated network density to compare changes in cohesiveness or saturation that relate to functional attributes, such as resilience^58^. We defined density as the number of links observed divided by the maximum number of possible links^58^, representing the probability that any given tie between two random nodes is present^59^.

We simulated two contrasting scenarios to investigate the potential effect of climate change on larval connectivity due to reduction of PLD with increased temperatures and the reduction of recruitment habitat due to climate change, since both could significantly alter metapopulation dynamics^21, 27^. In the first or “Current” scenario, we downscaled the larval connectivity matrices to the polygon unit (following the approach described by Alvarez-Romero, et al. ^21^) based on two factors: probability of connections between two polygons according to the connectivity matrix based on the PLD reported for each species in the literature, and the total area with giant kelp found within each polygon. The second or “Future” scenario employed shortened connectivity matrices due to warming and consequent restriction of giant kelp to highly persistent habitats, defined as potential climate refugia^20^. We calculated the reduction in PLD in fish and invertebrates following a 3°C increment in SST using previously described methods^21, 60^ (Supplementary Table 4). This increase in temperature is extreme, yet not unlikely for the end of the 21^st^ century under the highest IPCC Shared Socio-Economic Pathway (SSP5-8.5)^61^.

### Simulating recovery time for focal species

To project the expected effectiveness of transboundary marine reserve networks for recovery of exploited populations, in the absence of climate-change impacts, we simulated the effect of marine reserves on six focal species within the region (sea bass, whitefish, lobster, abalone, sea cucumber and sea urchin; Supplementary Table 1) using a deterministic, discrete-time logistic growth model with spatially implicit reserve and fishing zones:

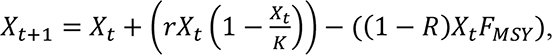

where *X_t_* represents biomass at time *t*, *r* is the intrinsic growth rate, and *K* is the carrying capacity. The last term represents harvesting of biomass outside the reserve, where the (1 – *R*) parameterization corresponds to the portion of biomass outside the reserve. We test three different scenarios of reserve coverage: *R* = (10%, 30%, 100%). The first two scenarios are representative of commonly cited protection targets, while the third scenario provides an upper bound of maximum attainable protection. We estimated population parameters and fishing mortality (*F_MSY_*) by applying a catch-only data-limited stock assessment method^62^ to catch data from 2000-2015 in Baja California, from Comisión Nacional de Acuacultura y Pesca (CONAPESCA) (Supplementary Table 5, Supplementary Fig. 4). We ran all simulations for 50 years, with initial biomass set at 20% of carrying capacity and fishing mortality outside the reserve held constant at *F_MSY_*. We considered a population “recovered” when the population size was within 90% of the theoretical equilibrium size (*X̄*):

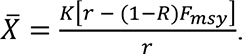

### Considering the impact of marine heatwaves on focal species biomass and recovery

For a subset of focal species of invertebrates with limited movement (abalone, sea cucumber and sea urchin; Supplementary Table 1, 6) vulnerable to marine heatwaves, we explored recovery for three climate-change scenarios by running a stochastic version of the model. We simulated the impact on the biomass of species based on the probability of a year experiencing marine heatwaves with a cumulative intensity at least as strong as those that impacted the California Bight in 2014-2015^34, 36–38^. We used 17 years of giant kelp forest community data (1999-2015), which integrate four different monitoring programs for the California Bight^37^, to model the rate of change of the density of the focal species following the 2014-2015 extreme marine heatwaves (henceforth “marine heatwaves”). We excluded monitoring data north of latitude 33.8°N and west of longitude 118.7°W because this area is subject to colder average temperatures, forms a separate sub-bioregion^63^, and giant kelp forest communities there are less impacted by marine heatwaves^64^ (for more details, see Supplementary Methods).

We used 0.25° grid cell Optimum Interpolation Sea Surface Temperature data set^65^ to estimate the average annual cumulative intensities registered in 2014-2015 marine heatwaves, based on the climatology for 1983-2012. Then we estimated projected marine heatwaves for the 2020-2100 period for three climate scenarios generated under each of three SPPs^66^ ─ SSP1-2.6, SSP2-4.5, SSP5-8.5 ─ from a multi-model ensemble mean derived from 11 Earth-System models from the Coupled Model Intercomparison Project Phase 6 (for more details see Supplementary Methods).

We derived the probability of a marine heatwave year using a generalized linear model with a binomial link function, fitted to marine heatwave years for the three climate scenarios. Then we used the empirically derived density change effect of a marine heatwave on abalone, sea urchins, and sea cucumber to yield the model:

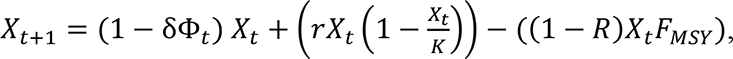

where *δ* ⋲ [0, 1] represents the density-reducing effect of the MHW, and *Φ_t_* = 1 if the year *t* has a MHW and *Φ_t_* = 0, otherwise. We ran 10,000 independent simulations of this model for each of the three species and three climate scenarios. All data analyses and simulations were conducted in R 4.0.4^67^.

### Mapping threats

We mapped climate and land-based threats that marine reserves cannot directly abate but can build resilience to (henceforth “threats”) at a 1-km^2^ grid resolution in the California Bight using the most comprehensive dataset available for the cumulative human impact (see Halpern, et al. ^68^). We classified those cells with high cumulative impacts (top 10%) for each sub-region as “highly threatened”.

For each of the three sub-regions in Baja California, we mapped two extractive activities that marine reserves can directly abate: small-scale commercial fishing and recreational fishing (henceforth “fishing”). We prioritized the analysis for these regions because they have less spatial information available. We mapped the total catch of small-scale fisheries (tonnes) at one-hectare grid resolution (Supplementary Table 7) inside authorized fishing polygons using 19 years (2000-2018) of catch information referenced to each fishing cooperative and individual permit holder (hereafter “concessions”) from CONAPESCA. Each fishery has a designated concession where cooperatives can extract specific resources (for more details see Supplementary Methods). Then, we estimated the probability of recreational fishing sites or banks in a 100-m^2^ grid resolution based on georeferenced commercial and non-commercial sport fishing maps (e.g., FISH.n.MAP CO., Baja California North, Sportfishing Atlas Baja California Edition) for the three sub-regions in Baja California (for more details see Supplementary Methods).

## Results

We provide 16 biophysical guidelines for designing transboundary networks of climate-smart marine reserves adapted for the California Bight and recommendations for their application (Table 1). The guidelines provide recommendations for habitat representation and replication (Supplementary Table 8), protecting critical and unique areas, incorporating connectivity, allowing time for recovery, and minimizing or avoiding local threats. Instead of addressing measures for adapting to changes in climate and ocean chemistry separately^9, 10^, we integrate climate-adaptation strategies within each of the other principles, to achieve a climate-smart network.

### Habitat mapping

We produced maps of the distribution of 31 coastal and island habitats from intertidal to deep-sea habitats for the four sub-regions (Fig. 2, Supplementary Table 3). Although southern California covers fewer degrees of latitude than northern and central Baja California, it represents almost half (∼46%) of the area of the California Bight. We mapped rocky intertidal and sandy beaches for ∼2700 km of coastal and island coastline. Our habitat mapping includes biogenic habitats such as eelgrass and surfgrass (*Phyllospadix* spp.) and different levels of giant kelp persistence. We also mapped subtidal sandy and rocky habitats at different depths and submarine canyons in all regions. We found no seamounts, guyots or other geomorphic features of importance for biodiversity in the territorial sea of the California Bight. We could not map surfgrass and rocky habitats deeper than 30 m for the three regions in Baja California. Finally, we found no giant kelp or estuarine habitats in Guadalupe Island.

**Fig. 2.**
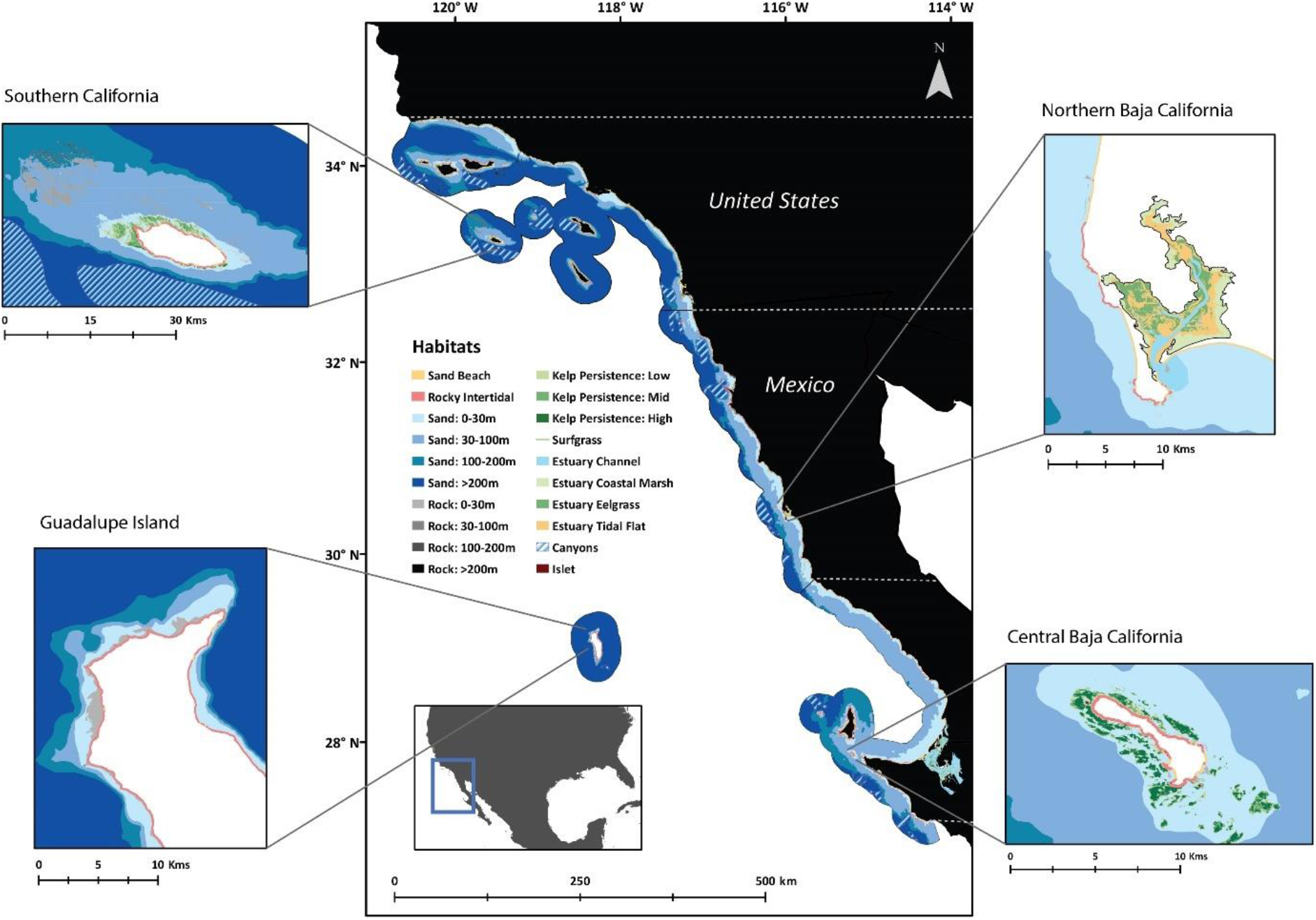
Distribution of marine habitats for the Southern California Bight. Inserts represent examples of intertidal, subtidal, estuary, and deep-sea habitats for each sub region. Southern California for San Nicolas Island with giant kelp forests, subtidal shallow and deeper habitats, northern Baja California for San Quintin Estuary, central Baja California for Isla Natividad intertidal and shallow habitats, and for Guadalupe Island intertidal and subtidal habitats at different depths. Dashed white lines represent the limits of each sub-region.

### Vulnerability of habitats and focal species to human threats

We received 55 responses to our expert-knowledge survey. Respondents ranked estuaries and rocky intertidal habitats as the most threatened habitats, whereas pelagic and deep-sea habitats ranked as the least threatened (Fig. 3a). Across habitats, marine heatwaves and resource extraction were identified as the most serious threats, while storm events and sea-level rise were the least threatening (Fig. 3b). Moreover, although the respondents ranked warming as the most threatening stressor, non-climate related threats, on average, scored higher (2.23 ± 0.17) than climate threats (1.94 ± 0.18).

**Fig 3.**
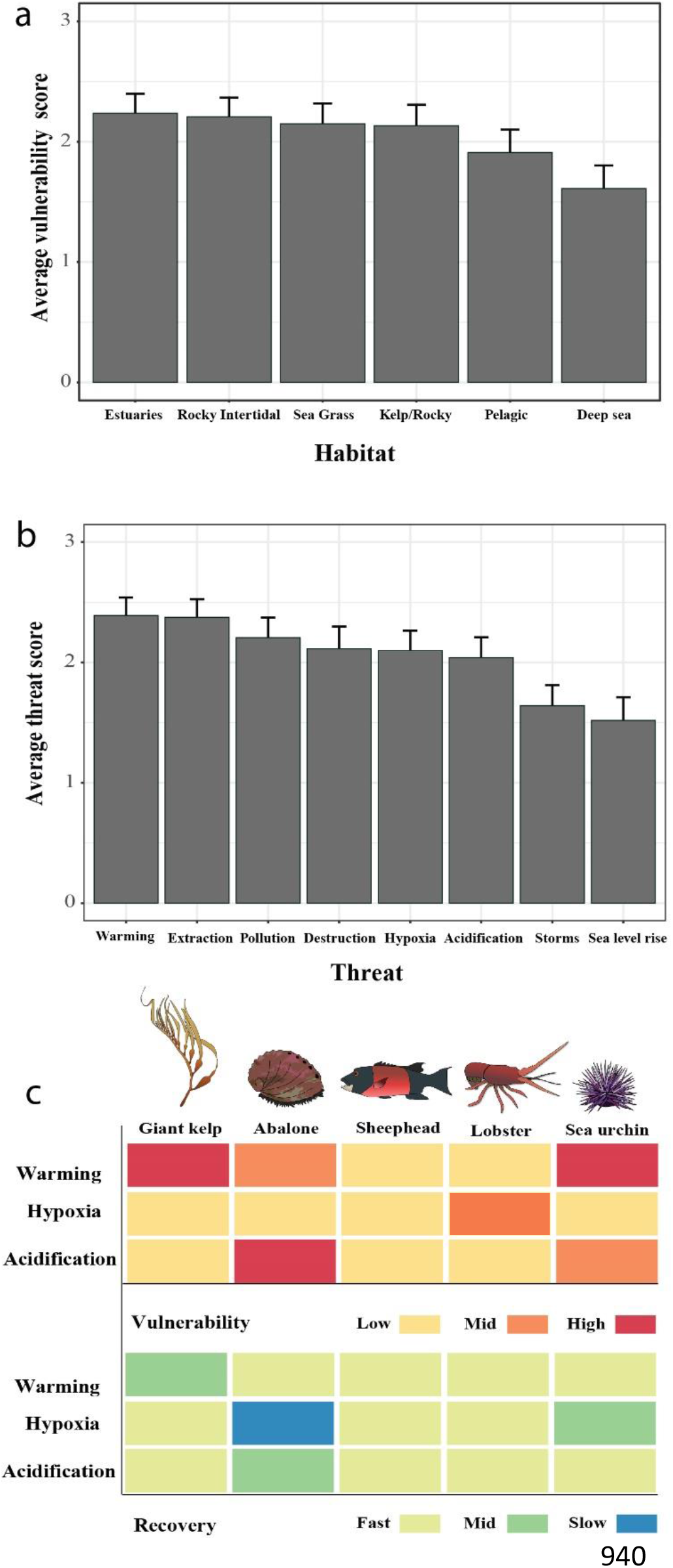
Vulnerability of habitats and focal species to human threats. Average score (± 95% confidence interval) for **a** vulnerability of marine habitats to human threats and **b** threat of individual stressors across marine ecosystems based on expert opinion surveys. **c** Ranking of focal species vulnerability to climate threats and their recovery rates based on scientific literature. Images credit: Katherine E. Dale.

Our risk assessment indicated that mobile species, such as California Sheephead and lobsters, are least vulnerable and recover fastest because they are least sensitive to acute climate stressors. In contrast, sessile species like giant kelp, or species with limited mobility, such as abalone and sea urchins, exhibit high and intermediate vulnerability to at least one climate stressor (Fig 3c). The recovery from climate stressors is intermediate for giant kelp, but slower for abalone and sea urchins. The combination of higher vulnerability and slow recovery renders these sessile or limited mobility benthic invertebrates most vulnerable to climate stressors (see Supplementary Results for detailed justification).

### Sea surface temperature variability as a proxy for climate refugia

Analysis of remote sensing SST and giant kelp canopy data revealed that high kelp persistence coincides with low temperature variability in southern California (p = 0.042), has no relation with temperature variability in northern Baja California (p = 0.745), and correlates with high temperature variability in central Baja California (p = 0.007) (Fig. 4). We also found that SST variability performs well compared to the observed high frequency bottom temperature variance (R^2^ = 0.87; Supplementary Fig. 5). Even when we excluded an outlier site (Punta Prieta) which has the highest temperature variability of all sites, we found a correlation coefficient R^2^ > 0.6. Our results suggest that in the absence of other metrics, SST variability may identify potential refugia for giant kelp and possibly kelp-associated species in the southern portion of the region (central Baja California).

**Fig 4.**
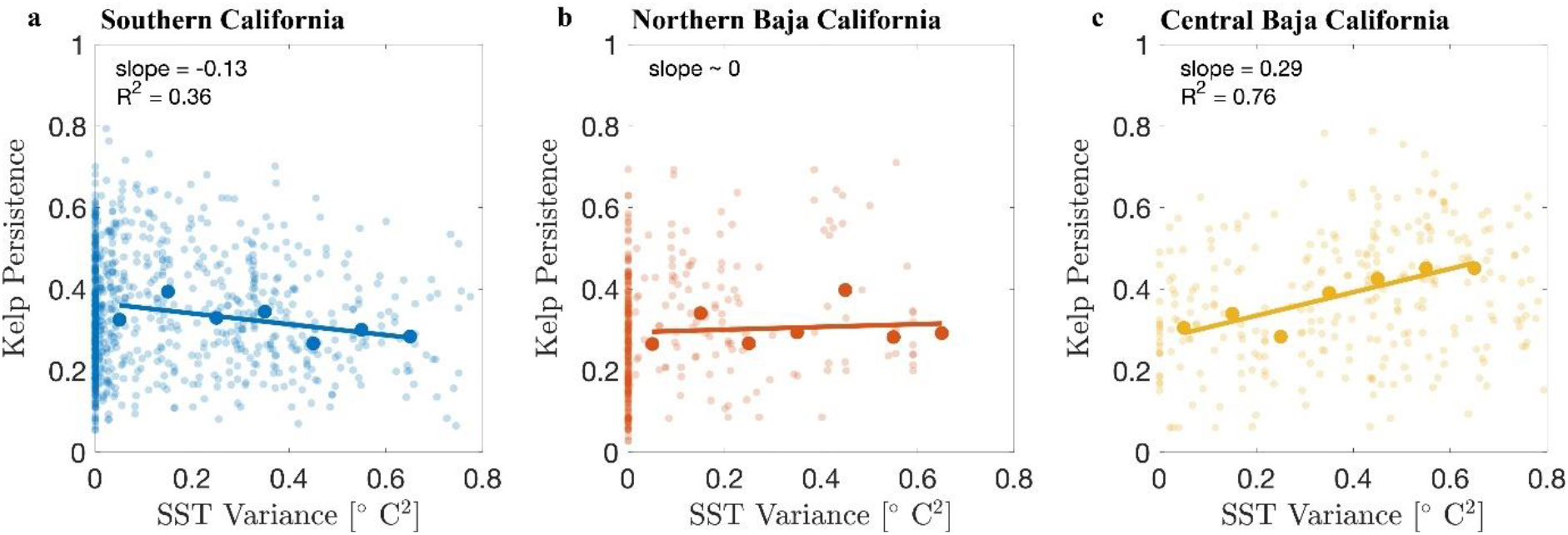
Sea surface temperature variability as a proxy for climate-change refugia. Relationship between giant kelp persistence and SST variability for **a** southern California, **b** northern Baja California, and **c** central Baja California. Raw data (light circles), binned-averaged values (large circles), and best fit to bin-averaged data (line) are shown.

### Transboundary movement of juveniles and adults

We found movement data for juvenile and adult individuals for seven shark, three bony fish, four seabird, four marine mammal, and one sea turtle species (Supplementary Table 9). Focal species are relatively large and migratory, with reported connectivity distances ranging from hundreds (e.g., white seabass, yellowtail) to thousands of kilometers (e.g., white sharks, blue whales; Supplementary Table 9) and temporal scales of two weeks (e.g., bottlenose dolphin, Black-vented shearwater) to over two years (white seabass). Greater connectivity was reported in northern Baja California, from north of Ensenada (31.8°N) and near the USA/Mexico border, followed by Punta Colonet (30.9°N), and areas in central Baja California, with important locations near the southern limit of the California Bight in Bahía Asunción (27.1°N). We also found areas of high connectivity between southern California and Guadalupe Island (Fig. 5).

**Fig 5.**
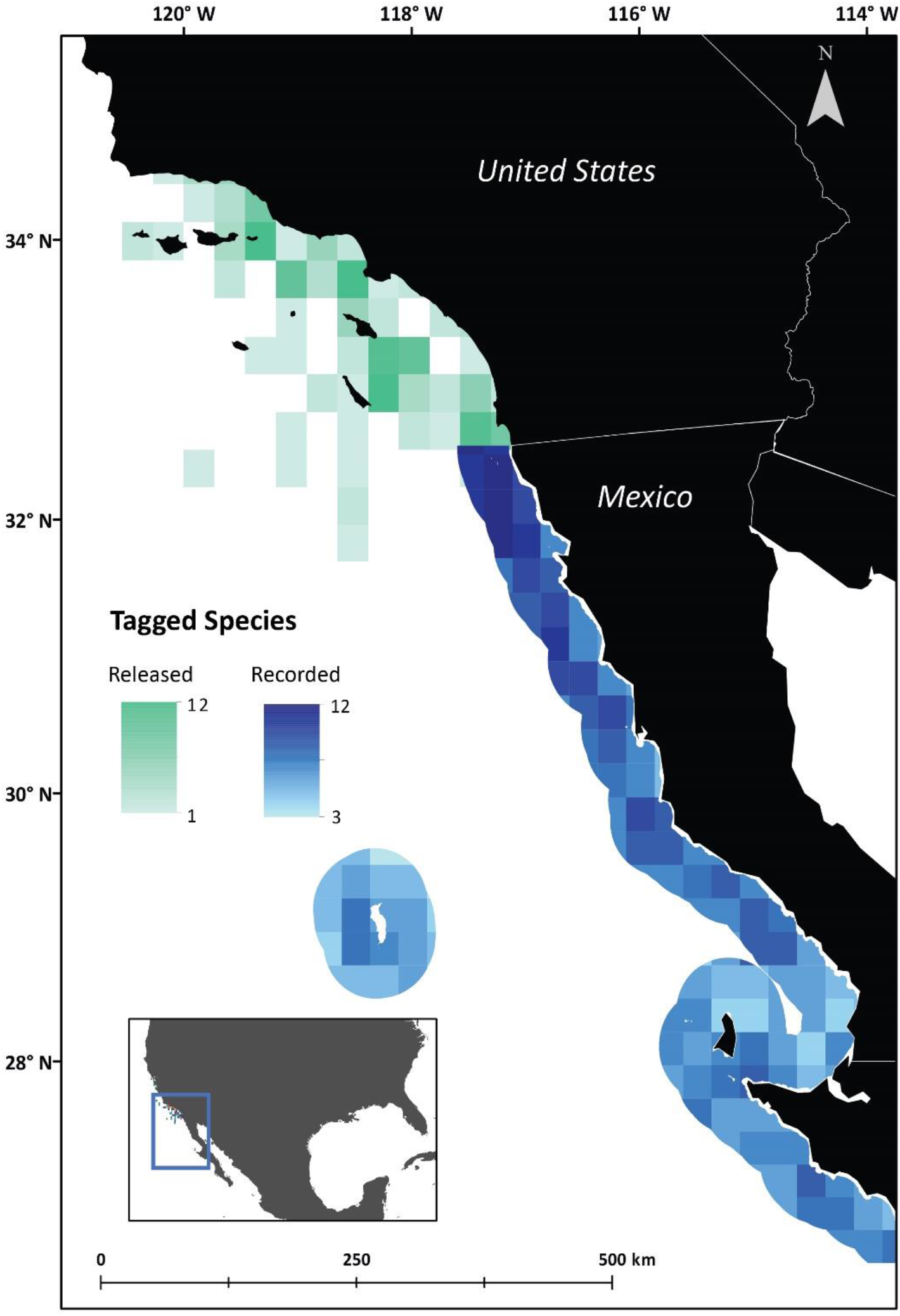
Transboundary connectivity of species moving from California to the Peninsula of Baja California. Green colors represent the number of studies that released species in southern California, and blue colors the number of studies that record species in Baja California. We consider blue quadrants with higher number of studies as quadrants with higher transboundary connectivity.

Tagging studies highlight transboundary nursery and foraging areas for white sharks in near-shore regions of central Baja California (Supplementary Table 9) and spawning areas for California Scorpionfish and Vermilion Rockfish in the USA-Mexico border. Breeding areas are also reported for Laysan albatross in Guadalupe Island, post-breeding dispersal areas for black-footed albatross offshore of San Quintín (30.5°N), in northern Baja California, and habitat use for the red knot in Guerrero Negro and Ojo de Liebre Lagoons (27.7°N) in central Baja California. In general, most studies did not report specific transboundary feeding, spawning, or foraging locations for most species, but rather broad general distributions along the California Bight (Supplementary Table 9).

### Transboundary larval dispersal model

Results of the larval dispersal models reveal that the California Current transports larvae southward throughout the year from California to Baja California, while a coastal undercurrent also transports larvae northward from Mexico towards California during summer and winter (Supplementary Movie 1). In the Current scenario, we observed transboundary larval connectivity along a stretch of coast covering ∼800 km, where larvae from Mexico travel ∼400 km north, reaching Point Conception (34.5°N) at the northern limit of the California Bight, while larvae from the USA travel ∼400 km south to El Rosario (29.8°N) in northern Baja California (Fig. 6a).

**Fig 6.**
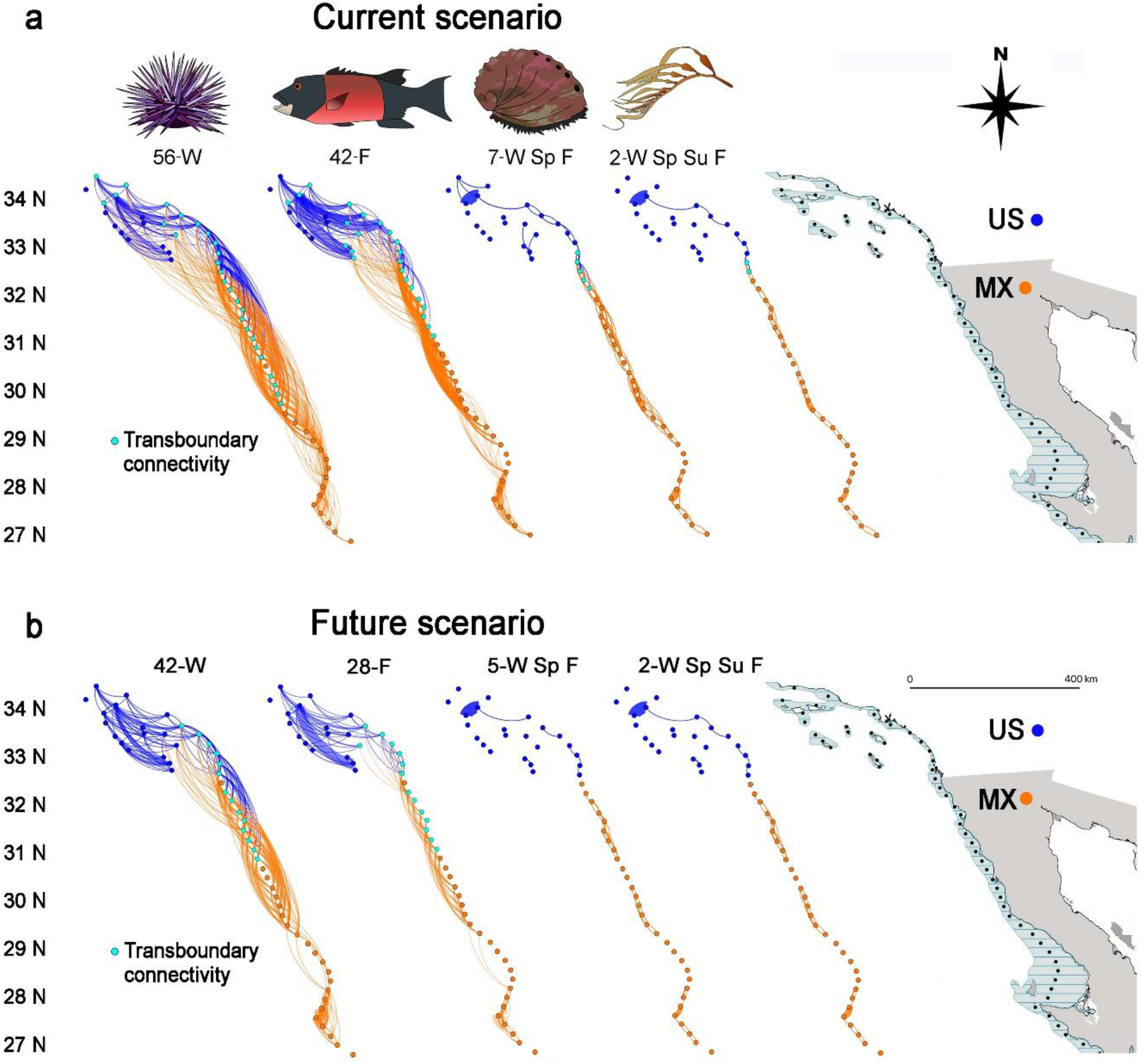
Networks of larval connectivity in the California Bight. Spatial networks of modelled larval dispersal for focal species (from left to right, sea urchin, California Sheephead, abalone, and giant kelp) between nodes delimited by the 200-m isobath. Connectivity polygons are shown on the far right. Line width represents the probability of larval dispersal (thicker lines have higher probability) and line colour the country of origin. Blue-coloured nodes and lines represent the USA and orange Mexico as larval origins, respectively. Sites involved in transboundary connectivity are highlighted in light blue color. For each focal species, we indicate the planktonic larval duration (PLD, in days), followed by a letter representing the spawning season (Spring = Sp, Summer = Su, Fall = F, Winter = W, See Supplementary Table 4). Images credit: Katherine E. Dale. **a** Current scenario, considering the PLD reported in the literature and the total area with giant kelp found within each polygon. **b** Future scenario, accounting for reduction of PLD and giant kelp habitat due to climate change. Sites involved in transboundary connectivity are highlighted in light blue color.

The strength of larval connectivity is a function of the geographic distance between sites, which generally decreases as the distance between sites increases. Connectivity patterns are strongly influenced by the life history of species, including planktonic larval duration (PLD) and spawning time. For example, propagules of the giant kelp (with a PLD of a few days to hours) move almost exclusively between adjacent sites on scales of < 25-50 km throughout the year, while larval dispersal of abalone (PLD = 7 days) occurs between sites 25-100 km apart, year-round. For these species, we found high levels of local larval retention (45% and 25% on average, respectively) compared to species with longer PLDs (average ≤ 7%). For all species, connections extend from southern California to central Baja California, but we found some isolated sites (with null probabilities of connectivity to other sites) for short PLD, such as in the offshore islands in southern California (henceforth “Channel Islands”) and areas in central Baja California.

In contrast, larvae of the California Sheephead (PLD = 42 days) can travel 75-500 km during the fall, and larvae of sea urchins (PLD = 56 days) can be transported 100-700 km during winter (Fig. 6a). Thus, transboundary connectivity is more important for species with longer PLD. The importance of transboundary connections varied by commercial species and country. For example, for sea urchin, 16% of all the larvae that settled within each country originated from the other country. On the other hand, for California Sheephead, 20% of all larvae in the USA came from Mexico and only 3% in Mexico from the USA. Although transboundary connectivity is more important for species with longer PLD in any single generation, long-term resilience over multiple generations depends on sites across the border that are tens to hundreds of kilometers away, even for species with short PLD.

Climate change is predicted to cause a significant decline in the number and strength of connections for all species. In the Future scenario, which considers the effects of climate change, the number of connections present and consequently the average density of larvae dispersing in the network decreased by half (range from −24% on giant kelp to −63% on abalone), the average probability of the connections reduced by an order of magnitude (range from −90.7% in California Sheephead to −96.0% in giant kelp), while local retention improved for all species except giant kelp (Supplementary Table 4). We found that binational connections were significantly reduced (61% and 52% lost for California Sheephead and sea urchins, respectively) or completely lost (giant kelp and abalone) either because larvae cannot reach as far or because stepping-stone connections disappeared due to the loss of giant kelp with low and intermediate persistence. For sea urchin and California Sheephead, the number of larvae crossing the border dropped to ≤ 3%. For species with short PLD, some sites become completely disconnected, forming independent sub-networks, or sites become only loosely connected through a few key sites, especially around the Channel Islands and in northern and central Baja California (Fig. 6b). Importantly, the Future scenario identified some sites (e.g., around the northern Channel Islands and Vizcaino Bay in latitudes 28-29°N) that may be crucial to avoid the collapse of connectivity in the region and should be prioritized for protection from additional impacts.

### Time for recovery of focal species and reserve coverage

The deterministic recovery model revealed that, overall, increased protection results in faster recovery and higher equilibrium biomasses. For all protection levels in the region (i.e., 10, 30, and 100%), sea cucumber showed the fastest recovery from fishing, at less than 10 years (Fig. 7a). Abalone and lobster showed the slowest rates of recovery, requiring between 31 and 47 years. Importantly, our results show that protecting 10, 20, or even 30% of abalone populations is not enough to allow recovery (Fig. 7b), highlighting that increasing reserve coverage and combining with other management actions is needed to facilitate population recoveries for slow-growing populations such as abalone. All fish and sea urchin species showed an intermediate recovery, requiring 15-20 years, regardless of reserve coverage. All populations reach recovery status fastest under a 100% protection, followed by 30% and then 10% (Fig. 7b).

**Fig 7.**
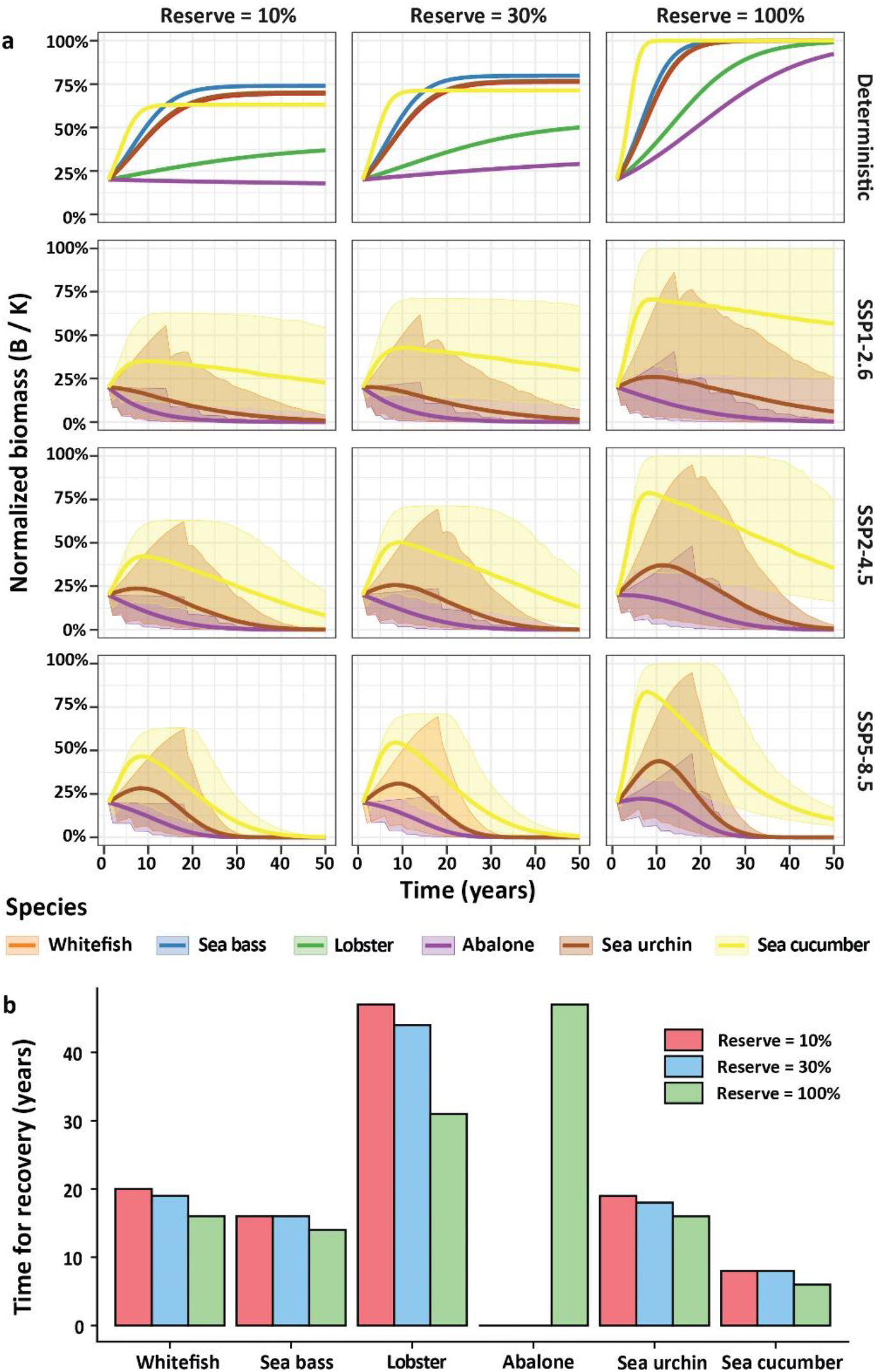
Simulated time to recovery for focal species under three protection scenarios in the California Bight. Recovery pathway of six species (mean ± standard deviation), **a** for scenarios of protection (10, 30, or 100%) and models. The first row of panel shows the deterministic model for all six species. Rows 2-4 show results from the stochastic models for vulnerable species (abalone, sea urchin, sea cucumber) to marine heatwaves under three climate scenarios (SSP1-2.6, SSP2-4.5, SSP5-8.5). **b** Time to reach 90% of equilibrium biomass for each of the seven species across three protection scenarios. Missing bars indicate no recovery within the simulated 50 years.

### Considering impacts of marine heatwaves on focal species recovery

We found an average cumulative marine heatwave intensity for 2014 and 2015 of 465.6 °C days and 684.5 °C days, respectively, for the nine pixels with monitoring data. When modelling future marine heatwaves, we found that by 2100 the probability of any given year experiencing a marine heatwave of this magnitude will be 0.46, 0.88, and 0.99 for scenarios SSP1-2.6, SSP2-4.5 and SSP5-8.5, respectively (Fig. 8). Importantly, if greenhouse gas emissions are not mitigated, the California Bight could be in a permanently extreme marine heatwave within the next 50 years (Fig. 8).

**Fig. 8.**
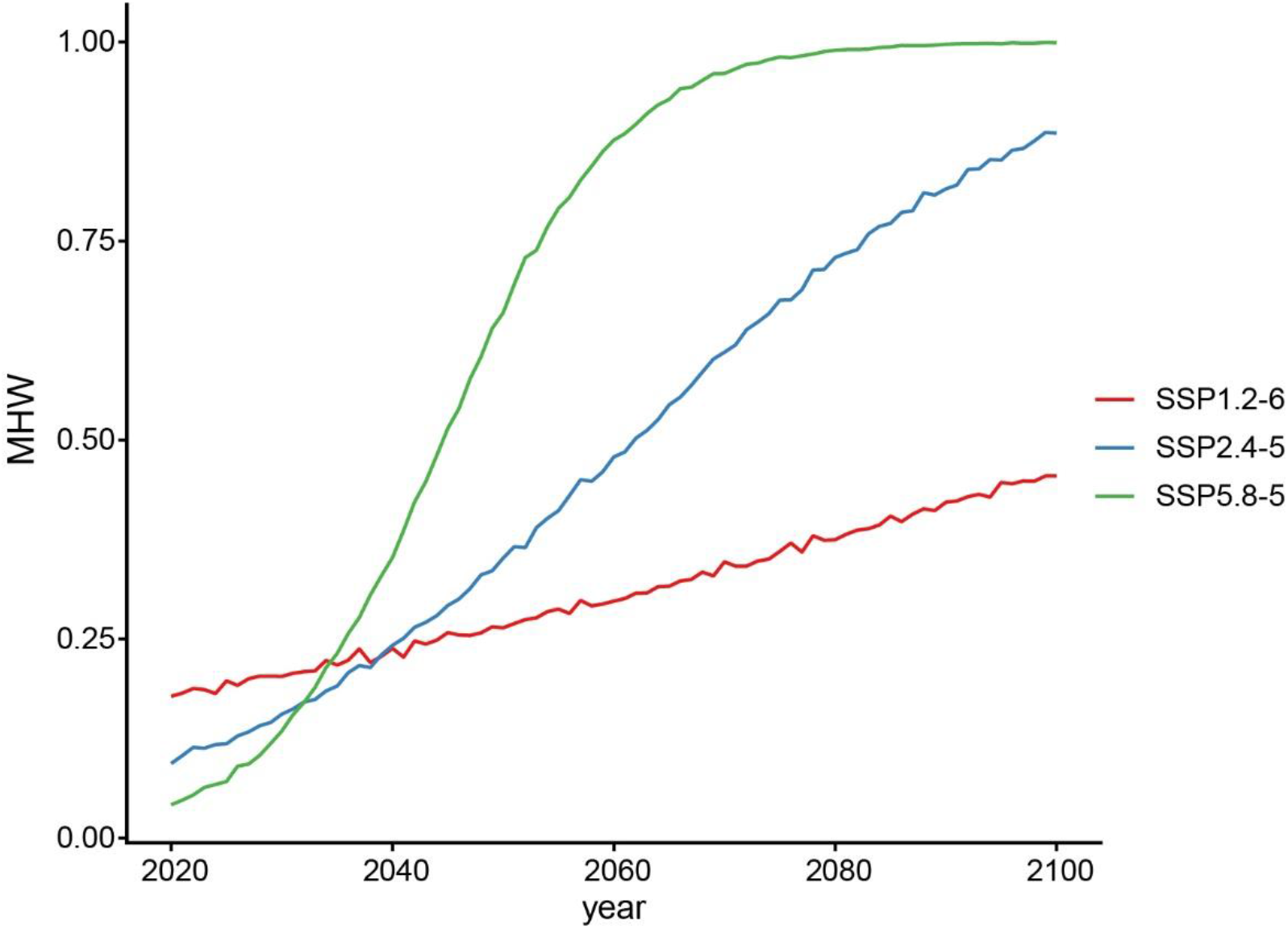
Probability of projected marine heatwaves for 2020-2100. Marine heatwaves under climate scenarios SSP1-2.6, 2-4.5, and 5-8.5 that match or exceed the cumulative intensities registered in 2014-2015 in the California Bight.

In linear models of yearly density change, we found significant differences between years with (2014-2015) and without (1999-2013) marine heatwaves for abalone, sea urchin, and sea cucumber (Supplementary Table 10; p < 0.01). Simulated yearly density of abalone, sea urchin, and sea cucumber species decreased during marine heatwaves (2014-2015) by 59.1%, 67.3%, and 72.4%, respectively.

When accounting for potential impacts due to marine heatwaves for vulnerable species, we found that no species reaches recovery status under any combination of emission scenario and reserve coverage, with abalone being particularly vulnerable (Fig. 7a). While not reaching equilibrium, sea cucumbers show the largest population sizes across climate and protection scenarios, but there was great uncertainty about those changes. Our simulations suggest that even rapidly growing species such as sea cucumber may not reach equilibrium biomass within the next 50 years even under 100% protection (Fig. 7a).

### Mapping threats

We found highly threatened areas that require management to support resilience, mainly near cities in southern California and northern Baja California (e.g., Los Angeles, San Diego, Tijuana, and Ensenada; Fig. 9a). However, in central Baja California and Guadalupe Island, we found highly threatened areas also in remote offshore sites. Both regions are isolated and scarcely populated, with Guadalupe Island located ∼250 km from the mainland of Baja California Peninsula. We also found an overlap of highly threatened areas with areas of high fishing pressure in northern Baja California but also in less populated areas further south at San Quintín (mainly recreational fishing) and El Rosario (primarily commercial fishing) (Fig. 9b-c). On the other hand, fishing pressure in central Baja California was lower and mainly concentrated near Punta Eugenia and Bahía Tortugas for commercial fishing and south of Cedros Island for recreational fishing. Sea urchin and sea cucumber are the most important commercial fishing catch in northern Baja California, lobster and abalone in central Baja California, and abalone in Isla Guadalupe (Supplementary Table 7).

**Fig. 9.**
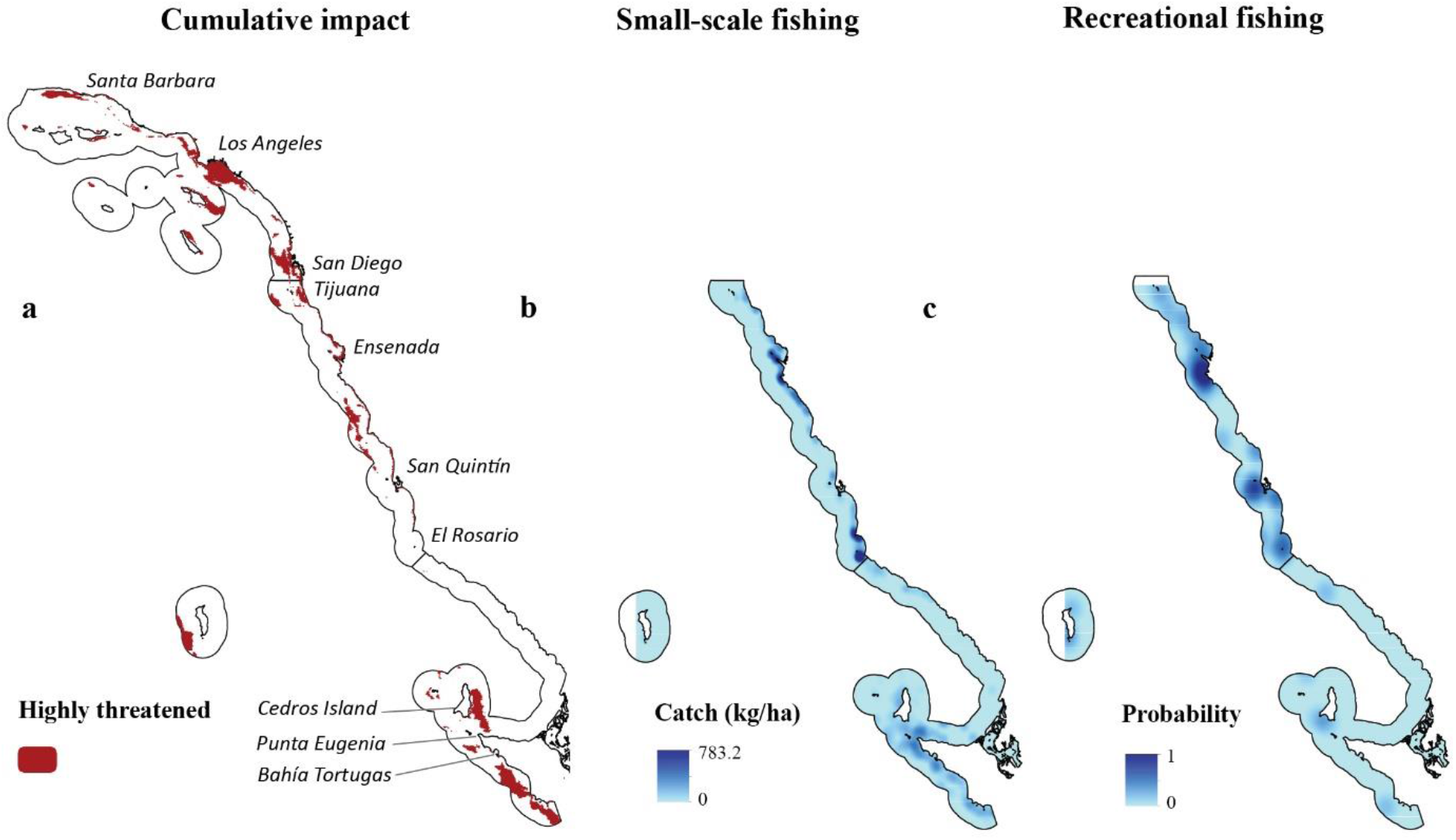
Distribution of threats and fishing in the California Bight. Maps of **a** highly threatened areas (top 10% of cumulative impacts of 13 threats), **b** cumulative catch for six small-scale fisheries and **c** probability of recreational fishing in a 100-m grid square in the three regions of Baja California.

## Discussion

Our results show that although southern California occupies ∼25% of the California Bight extension, spanning the USA-Mexico border, it contains almost half of the marine habitats and supports strong ecological connections with northern Baja California. This transboundary connectivity is already significant for populations with long planktonic larval duration (PLD) and highly mobile adults and juveniles, and is likely to become more important as species shift their distributions in response to changing environmental conditions. Given that many populations of ecologically and commercially important species depend on both countries, it is essential to protect important transboundary areas for migratory species and larval dispersal to recover and maintain populations, communities, and genetic flows^10, 69^ and benefit both conservation and fisheries^47, 50, 70, 71^.

Under a future climate scenario, we found that many areas in the California Bight become isolated and binational connections will diminish or be lost, especially for species with short PLD. However, local retention improves for most species, suggesting that establishing large marine reserves in areas that will become more isolated is critical to maintaining self-replenishment and supporting local populations^61, 72^. Like other studies^21, 27^, we found that the strength of connections weakens, the overall larval recruitment decreases, and that some nodes for species with short PLD may become disconnected. Under this future climate scenario, networks of marine reserves will need to prioritize the protection of key stepping-stone nodes to avoid the collapse and fragmentation of larval dispersal in the region. If not adequately protected from fishing, depleted populations’ limited supply of larvae could lead to genetic bottlenecks^73^ and local population collapse^74^, with economic implications.

Our results indicate that SST variability is a good proxy for climate refugia, particularly for central Baja California. Areas with high-frequency daily variability in ocean conditions (24 hours or less) can provide refuge^30, 75, 76^, similarly to deeper nearby habitats where vulnerable sessile species can survive adverse conditions and mobile species can retreat^77, 78^. The observed lack of relationship between SST variability and giant kelp persistence in southern California and northern Baja California may imply that giant kelp there are less limited by nutrient availability and temperature extremes than giant kelp near their southern distribution limit in central Baja California^79^. Therefore, it is a research priority to assess whether highly persistent giant kelp provide refuge for vulnerable species to climate impacts.

The probability that the California Bight will be subject to similar marine heatwaves as extreme as those in 2014-2015 in the following decades is high, and it becomes more likely every year while carbon emissions continue to rise, consistent with global analysis^80^. Even if carbon emissions can be reduced in the next decades, the California Bight will most likely face new extreme events. Along with marine heatwaves, resource extractions are the main threats in the region, which can have interactive and cumulative effects and degrade the resilience of marine ecosystems^81^. Notably, when we include the potential impacts of future marine heatwaves on the recovery of vulnerable species (sessile or limited mobility), our modelling results suggest that even with high levels of protection, these species of commercial and ecological importance will not fully recover in subsequent decades. Results from the expert-opinion survey and literature review corroborate the notion that species with limited movement and with slow recovery (e.g., abalone) are more vulnerable than mobile, fast-recovering species. Protecting climate refugia from extractive activities might be the best available climate-adaptation strategy^11^ to buffer the impacts of future marine heatwaves on vulnerable species.

Coordinated conservation efforts for entire ecoregions will support climate-smart designs for biodiversity conservation and fisheries management to a greater degree than networks developed separately by each nation^50, 71, 82, 83^. We linked climate-smart strategies^11, 17^ (e.g., assessing climate impacts and vulnerabilities, representing climate refugia, incorporating connectivity, forecasting effectiveness of protection) using kelp forest ecosystems and associated focal species considering future threats from climate change. Notably, the climate-smart focus of our work addresses the growing need to meet post-2020 conservation targets and protect 30% of the oceans by 2030 while adapting to climate change^23, 24^.

We developed biophysical guidelines to ensure that a transboundary network of marine reserves in the California Bight would complement the existing network in California^63^, by adapting and refining existing best practice guidelines from California, Mexico, and other regions^9, 10, 40^. Setting representation targets at 30% of the distribution of each habitat in each biogeographic sub-region will protect ecological communities and processes in the California Bight^9, 10, 22^ and support adequate recovery for a range of focal species^84^. However, we found certain habitats (e.g., seagrass, giant kelp forests) to be more vulnerable to threats and may require higher levels of protection^49^. Habitat representation targets can be adjusted if robust fisheries measures are in place^9, 10^, but will need to consider the between-country differences in catch and management policies^53, 85^.

Many focal migratory species in the California Bight can move for hundreds to thousands of kilometres^85^. Protecting these species requires a combination of marine reserves and other management tools, such as gear restrictions, catch quotas, seasonal bans, or species moratoria, among other measures^9, 70^. Although marine reserves cannot fully protect species with long-distance (>10s km) movements that migrate outside their borders, they can protect areas critical for their life history (e.g., nursery, foraging, spawning)^9, 10, 70^. For these actions to effectively be in place, protecting critical areas across the border will require increased investment in international cooperation and governance^86^. Moreover, we found little information about critical transboundary sites, highlighting the need to develop research programs and collaborations to generate this information. Notably, setting adequate protection and management strategies will also require a better understanding of the populations of focal migratory species within and across borders in the California Bight^85^.

Our findings that transboundary connectivity is more important for species with longer PLD agree with empirical data on the genetic structure of these species. For example, studies found strong genetic differentiation for neutral genetic markers for species with short dispersal distances, such as giant kelp^87^ and pink abalone^73^, between southern California and populations from Baja California. In contrast, studies found lack of genetic structure for California Sheephead between California and Baja California^88^ and purple and red sea urchin from Washington to southern California^89, 90^. To meet international commitments^24^, fully protected areas will need expansion in California (currently ∼10%) and in Baja California (currently < 1%). This provides an opportunity for both regions to coordinate efforts and maximize conservation and fisheries benefits based on biophysical principles and modelling of larval dispersal.

We recommend establishing marine reserves in areas where threats can be managed effectively within reserves in the California Bight. Areas with high cumulative impacts (e.g., coastal development, pollution, runoffs) are likely degrading ecosystem health, fisheries productivity, and resilience to climate change (reviewed by Green, et al. ^9^), preventing marine reserves from producing the expected benefits^9, 10^. However, these are general recommendations since reducing overfishing inside marine reserves, combined with restoration actions and other management strategies that directly address those threats, can build resilience^3, 5, 7^ to threats not directly managed by marine reserves and contribute to the recovery of degraded areas. Therefore, the decision to protect or restore highly threatened areas requires cost-benefit analysis on a site-specific basis^91^ and other considerations such as ecological connectivity.

Prior to our effort, there were no comprehensive high-resolution maps of small-scale fisheries catch in the Peninsula of Baja California, limiting our capacity to conduct cost-effective marine spatial planning. Although information on recreational fishing existed^53^, we updated and extended the mapping to cover the three sub-regions in Baja California. These maps, combined with existing data from socioeconomic activities from California, provides the unique opportunity to guide marine spatial planning and identify priority areas for conservation that meet our biophysical design principles while minimizing the potential conflict from implementation to stakeholders’ activities^48^.

We note that our findings are subject to some caveats. We simulated passive larval dispersal and did not consider other biological traits (e.g., larval mortality, behavior, and settlement competency)^92^. However, we did tailor our model to the availability of giant kelp. Successful recruitment of larvae is linked to the quality and suitability of available habitats^22, 93, 94^. Although our climate scenario is a realistic expectation if carbon emissions are not mitigated in the 21^st^ century^61^ (decrease in PLD following a 3°C warming tailored to the availability of only highly persistent giant kelp habitat), it is an extreme scenario. Thus, if our model was to be used for identifying climate-smart networks of marine reserves, we recommend adjusting the model to consider multiple emission scenarios (e.g., SSP1-2.6, SSP2-4.5, SSP5-8.5) based on projected multi-model ensemble means for the California Bight.

While it is currently unfeasible to obtain information for all habitats and species in a region, we chose giant kelp forests because of existing information on their persistence^20^ and the availability of long-term information for associated focal species^37^. Also, kelp forests are threatened by marine heatwaves globally^35, 95^, acting as early indicators of climate-change impacts to other ecosystems. However, as more information becomes available, similar assessments for other vulnerable ecosystems, such estuaries, and rocky intertidal habitats should be conducted. Although our analysis found no positive correlation between SST variability and persistence of giant kelp forests for southern California and northern Baja California, SST variability may still be a good proxy for climate refugia for these regions, but anthropogenic activities may be eroding the persistence of giant kelp forests and masking the potential relationship. For example, many of the kelp forests in mainland southern California and northern Baja California experience high human impacts (e.g., run-offs, overfishing), while central Baja California is sparsely populated and less impacted. Also, our analysis compared SST variability with giant kelp persistence, and we did not include other species vulnerable to climate change such as sessile invertebrates. Therefore, it is a research priority to understand the drivers and synergies controlling species and habitat resilience to extreme events.

We also acknowledge caveats in our recovery model which did not include the potential benefit of larval dispersal from climate refugia, where populations may be less impacted by marine heatwaves. Also, we simulated the loss of biomass based on empirical data of the impacts of the 2014-2015 marine heatwaves from giant kelp forest monitoring surveys^37^ at shallow depths (typically less than 15 m). Thus, our results need to be taken with due care, as some deeper populations of invertebrate and sessile (or limited mobility) species might be less impacted^77, 78^, survive, and support the replenishment of nearby affected areas. Finally, we used catch-only methods to estimate population parameters and fishing mortality. Recent work has shown that these methods may produce biased estimates^96^, however our parameter estimates are similar to those reported in other studies^97^. Regardless of the potential limitations, our results are consistent with other work that suggest the need to increase the coverage of marine reserves to rebuild marine life, including protection of climate-refugia^5, 98–100^. Notably, other management actions such as fishing moratoria, catch quotas, and repopulation of vulnerable species will be needed to complement marine reserves.

Fully protecting 30%^24^ of the California Bight by 2030 will require national and transboundary policies and political will. Importantly, there is a need to integrate biophysical guidelines with socio-economic and governance principles to produce effective, equitable and robust policies and practices^42, 101^ while considering cultural and management differences across the border. Unfortunately, despite the scientific capacity and established collaboration among institutions and research groups between the USA and Mexico^31^, existing political cooperation matches neither the level of ecological connectivity observed^85^, nor the needs identified under current and projected climate impacts. Urgent, coordinated binational action needs to be taken to preserve fisheries and conserve biodiversity in the region. It is a grand challenge, given the strong asymmetries in economic wellbeing, governance, implementation capacity, resources, and language, among other barriers.

Marine reserves in Baja California will require co-management that includes local fishing cooperatives, complemented with other effective management strategies^42^. Some well-managed fishing concessions may need less protection, and in some cases, coordination with improved management and restoration actions may achieve biodiversity, fisheries, and climate-adaptation objectives. On the other hand, California has the legislative infrastructure for expanding its existing network of marine reserves and the experience in marine spatial planning to create synergies across the border. This collaborative and socio-ecological setting creates a unique opportunity for the California Bight to implement transboundary and climate-smart marine spatial planning and influence marine conservation worldwide.

Here, we provide a case study that links biophysical design principles for transboundary climate-smart networks of marine reserves. Our analysis suggests that achieving climate-smart status requires integrating multiple adaptation strategies such as protecting climate refugia and considering the implications of climate change for ecological connectivity and protection efficiency. Given that many marine ecoregions worldwide are shared by nations^43^, our biophysical guidelines and recommendations can inform other regions’ aspirations to achieve post-2020^23, 24^ protection targets. These regions will need to develop biophysical dispersal models to understand patterns of connectivity, identify potential climate refugia and levels of protection needed to maximize biodiversity, fisheries, and climate adaptation outcomes. Notably, to design climate-smart networks of marine reserves, they will need to coordinate research programs and policies while considering cultural, governance, and management differences across borders.

## Acknowledgements

N.A.-D. acknowledges support from the Fundación Bancaria ‘la Caixa’ under the Postgraduate Fellowship (LCF/BQ/AA16/11580053), from the University of Queensland under the Research Training Scholarship, and from the Estate Winifred Violet Scott for a research grant. O.A-O acknowledges the UC-Mexus Collaborative Grant (2016: CN-17-133) received for first workshop. F.M, A.S, C.B.W, MP, A.M.-V and S.F acknowledge the support of the US NSF (grants BioOce 1736830 and DISES 2108566). We are deeply grateful to all participants of the workshops.

