## Supplementary Material for "Towards transboundary networks of climate-smart marine reserves in the Southern California Bight"

### Supplementary Methods

#### Vulnerability of habitats and focal species to human threats

*Expert-knowledge survey*

In June-July 2020, we contacted experts in the California Bight to quantify the vulnerability of six marine habitat types (estuaries, rocky intertidal, seagrass, kelp forest/rocky reefs, deep sea and pelagic) to eight major threats (marine heatwaves, ocean acidification, hypoxia, sea-level rise, storms, resource extraction, pollution, and physical habitat destruction). Experts represented knowledge of six major topics: fishing and local ecological knowledge, ecology, oceanography, fisheries management, aquaculture, and conservation/restoration. These experts were identified by compiling the suggestions of working group members. We asked the experts to complete the survey, focusing on the region and ecosystem type for which they were considered an expert.

*Literature review*

Our search terms included "climate change", "warming", "heatwave", "temperature increase", "acidification", and "hypoxia" as key words. We defined the recovery rate as the number of years a particular species would require recovering its functional role within the system after exposure to a given stressor. For instances lacking empirical studies (e.g., effects of hypoxia or ocean acidification on California Sheephead (*Semicossyphus pulcher*)), we used literature from related taxonomic groups, as well as expert knowledge to score vulnerability and recovery rate.

#### ROMS implementation

The ROMS model solves the dynamic ocean equations based on finite differences[1]. We implemented the model in the northeast Pacific Ocean from Oregon (44°N), USA, to Chiapas, Mexico (14°N) (Figure S4.2). The model's spatial resolution varies from ~7 km near the coast to 20 km at the open western and southern boundaries. It has 40 sigma levels along the water column following the stretching parameters $\theta S = 6.0$, $\theta b = 0.2$ y $hc = 10$ m. It includes climatology (calculated based on the 2000-2020 period) and heat and salt fluxes derived from the Comprehensive *Ocean*-Atmosphere Data Set as atmospheric forcing[2]. We used the Advanced Very High-Resolution Radiometer[3] data set as the reference surface temperature to calculate thermal relaxation forcing at the surface, which is proportional to the temperature difference between the model and a reference surface temperature.

We obtained the bathymetry from ETOPO2[4] and used monthly average for the conditions for the open boundaries from the Simple Ocean Data Assimilation[5]. We parameterized turbulent mixing processes smaller than the grid-scale using a k-profile[6]. We calculated wind forcing using 6-hour average wind stress calculated from ERA-Interim at 10 m above sea level. We used the ROMSTOOLS subroutines[7] to fit the forcing fields, bathymetry, and open boundary conditions. We spun the model up for ten years using the climatology fields and then ran it on for 20 years.

**Mapping small-scale fisheries catch**

We randomized locations for catches of each species within each concession. Maps for California Sheephead and kelp sea bass (*Paralabrax clathratus*) were generated by selecting giant kelp forests (*Macrocystis pyrifera)* near each concession landing site and randomizing locations within kelp forest edges for California Sheephead or core for sea bass, following previous approaches[8]. Lobster concessions were delimited by kelp cover and depth (traps are not commonly set inside giant kelp forests or beyond 300 m depth) while abalone, red sea urchin, and sea cucumber concessions were delimited by diving depth (from 5-30 m) following published methods[9]. Data catch records from CONAPESCA includes date, catch in kg, landing site, and number of vessels used for fishing. California Sheephead and kelp seabass are mainly caught by hook and line, while spiny lobster (*Panulirus interruptus*) is caught with baited traps, and abalone (*Haliotis* spp.), red sea urchin (*Mesocentrotus franciscanus*), and sea cucumber (*Apostichopus* spp. and *Parastichopus* spp.) are hand-picked by hookah divers. California Sheephead and kelp seabass are fished under “finfish” commercial permits, mainly with hook and line onboard artisanal vessels, with a small finfish mid-range fleet. Spiny lobster, abalone, sea urchin, and sea cucumber are fished under commercial fishing permits or concessions only with authorized traps and diving gear within the authorized fishing area.

California Sheephead and kelp seabass catch data were geo-reference assuming a higher abundance of California Sheephead in the outer zones of the kelp forests, while kelp seabass inhabits middle and outer zones of the forest[8]. Based on catch records landing sites and species, we assigned catch a random location either outer or in the core zones of the kelp forests surrounding landing sites. Spiny lobster catches were geo-referenced in each authorized fishing permit or concession. Given that traps are not placed inside kelp forests nor dropped deep, we excluded areas covered by kelp and depths greater than 300 m. For abalone, sea urchin, and sea cucumber catch we geo-reference each fisheries landings using a randomizing function within each authorized fishing permit or concession in depths between 5-30 m. It is important to notice, the specific point locations generated do not account for actual fishing sites or actual hotspots within each area.

##### Mapping recreational fishing

We calculated the kernel density of recreational fishing sites using a 100-$m^2$ grid to assign a higher weight to areas closer to each other. We also consulted existing documents such as "fishing site Atlas" from CONAPESCA to corroborate the geographic position of fishing sites with duplicate names or different nomenclature. These fishing documents contain latitude and longitude of all fishing sites.

#### Supplementary Results

##### Vulnerability and recovery of focal species to climate threats

We provide detailed justifications for the evaluation of the vulnerability and recovery of sheephead, lobster, abalone, sea urchin, and giant kelp to marine heatwaves, hypoxia events, and ocean acidification.

##### California Sheephead (*S. pulcher*)

| Threats | Vulnerability | Recovery |
|---|---|---|
| Marine Heatwaves | Low | Rapid |
| Hypoxic Events | Low | Rapid |
| Ocean Acidification | Low | Rapid |

##### Marine heatwaves

California Sheephead respond to the thermal gradient of the Southern California Bight and occur in small numbers in central and northern California because waters are too cool to sustain breeding populations[10]. There is no evidence of a warming impact, so sensitivity is coded as low and response as rapid.

#### Hypoxia

California Sheephead are highly mobile and can easily avoid hypoxic events, so their sensitivity is coded as low and their recovery as rapid.  There is no evidence of hypoxia impacts.

#### Ocean Acidification

California Sheephead are highly mobile and can easily avoid acute low pH events, so their sensitivity is coded as low and their recovery as rapid. There is no evidence of ocean acidification impacts.

#### Lobster (*P. interruptus*)

| Threat | Vulnerability | Recovery |
|---|---|---|
| Marine Heatwaves | Low | Rapid |
| Hypoxic Events | Medium | Rapid |
| Ocean Acidification | Low | Rapid |

##### Marine heatwaves

Lower temperatures and upwelling are linked with higher rates of spermatophore deposition[11]. Therefore, the sublethal impact of high temperatures will likely include a lower reproductive output. There is no evidence of lethal impacts at current conditions. Lobster mobility and wide range of experienced temperatures indicate that their response to marine heatwaves will be rapid.

##### Hypoxia

The mobility of lobsters will likely allow them to move away from hypoxic waters. This is corroborated by monitoring data showing that lobsters were unaffected by prolonged hypoxia at Isla Natividad, BCS[12].  However anecdotal evidence from the American lobster fishery in Cape Cod reveal that when mobility is restricted by traps, they may suffer mortality from a hypoxic event due to water column stratification. We decided on a sensitivity score of medium as there are lethal impacts at current conditions, but they only occur in specific cases. Response was coded as rapid since lobsters are mobile can avoid this threat.

##### Ocean Acidification

Much of the literature on lobsters and ocean acidification deals with the impact on the larval and juvenile stages, which finds that recruitment and settlement can be negatively impacted but the degree varies widely across species and lobster is not explicitly discussed. There is no evidence of lethal or sublethal impacts under current ocean acidification conditions. Moreover, meta-analysis of published studies found an overall non-significant or positive effect for crustaceans for several response variables considered, possibly due to their ability of controlling extracellular pH through active ion transport resulting in greater tolerance of acidification[13]. Based on this information, sensitivity is coded as low.

**Abalone (*Haliotis.* spp.)**

| Threat | Vulnerability | Recovery |
|---|---|---|
| Marine Heatwaves | Med | Rapid |
| Hypoxic Events | High | Slow |
| Ocean Acidification | High | Low |

**Marine heatwaves**

Experimental and monitoring studies in Isla Natividad[14] suggests warming less of an issue compared to low dissolved oxygen for growth. Juvenile abalone potentially acclimate to high temperatures and mortality was limited, but activity decreased. Recovery is likely faster if acclimation is possible.

**Hypoxia**

Mass die-offs likely caused by hypoxic events near Isla Natividad[15]. Growth is depressed[14] because hypoxia leads to mass die-offs and abalone are slow growing, recovery will likely be slow. A study measuring physiological responses found that juvenile green abalone (*H. fulgens)* can adapt to increased heat and hypoxia when events are independent. However, they are unable to adapt during combined warming and hypoxic events[16].

**Ocean Acidification**

Juvenile growth depressed for red abalone, which may lead to decreases in reproduction and increases in predation risk related to prey size[17]. The negative effects may negatively impact different larval stages[18]. Acidification was found to reduce fertilization success of red abalone, with high temperatures mitigating this negative impact[19]. Recovery likely to be moderate due to potential decreases in reproduction linked to depressed growth.

**Sea urchin (*Strongylocentrotus purpuratus* and *Mesocentrotus franciscanus*)**

| Threat | Vulnerability | Recovery |
|---|---|---|
| Temp Increase | High | Rapid |
| Hypoxic Events | Medium | Moderate |
| Ocean Acidification | Medium | Rapid |

**Marine heatwaves**

Mass die-off events have been witnessed at current thermal extremes. Studies describe lethal impacts of fertilized eggs at 25°C[20] which are temperatures reached in marine heatwave events in the region[21,22] so we characterize as a sublethal (to the adult form) impact. We are not aware of literature discussing functional recovery after lethal impacts of temperature, but anecdotal evidence of barrens eliminated by storms recruit new adult urchins within a few months. Therefore, coding recovery as rapid.

**Hypoxia**

Studies show that while lethal impacts are not likely in current or near-future exposures, there is a high likelihood of sublethal impacts, namely decreased grazing rates, for substantial periods of time under current conditions[23]. Further, recovery to their full functional role is estimated to be quite slow.

**Ocean Acidification**

Studies demonstrate a 24% reduction in fertilization success in near-future ocean acidification scenarios for the sea urchin *Heliocidaris erythrogramma*[24]. Studies also purport that adults can acclimate to elevate pCO2 but that negative impacts may be experienced by larvae and juveniles in the next generation[25]. Therefore, sensitivity is coded as medium due to the sublethal impacts at current conditions. However, there is no evidence to suggest a loss of functional role in the ecosystem, so recovery is coded as rapid.

**Giant Kelp (*M. pyrifera*)**

| Threat | Vulnerability | Recovery |
| --- | --- | --- |
| Marine Heatwaves | High | Moderate |
| Hypoxic Events | Low | Rapid |
| Ocean Acidification | Low | Rapid |

**Marine heatwaves**

Giant kelp along the Southern California Bight is sensitive to an absolute thermal maximum that results in a rapid loss; therefore, their sensitivity is characterized as high. In terms of recovery, studies found that kelp resilience is not correlated with sea surface temperature but after a heatwave, most kelp forests re-established within one generation[22,26] indicating a Rapid recovery. However, other studies found longer times for recovering and giant kelp forests in their southern limit distribution are more vulnerable and might be slower to recover in Bahia Asuncion and Punta Prieta in central Baja California following marine heatwaves[27-30].

**Hypoxia**

The literature is inconclusive regarding the impacts of hypoxic events on giant kelp. Crowder, et al. [31] explain that there may be a conflicting impact since low levels of oxygen may enhance photosynthesis but impair respiration. The best available science lacks evidence of any specific sublethal or lethal response of kelp to current hypoxic events, therefore, they were scored Low for sensitivity and Rapid for their response to this climate stressor.

**Ocean Acidification**

Much of the literature related to the relationship between kelp forests and ocean acidification is either focused on early life history stages or the ability of kelp forests to ameliorate low pH water chemistry.

#### Supplementary Tables

**Supplementary Table 1.** Focal species used for each analysis. Note that some focal species may use information for more than one species in that group (e.g., abalone, see Supplementary Results and Table S4.4-6, S4.10).

| **Focal Species** | **Vulnerability and recovery rate** | **Larval dispersal** | **Recovery time** | **Recovery time with marine heatwaves** | **Small-scale fishing** |
|---|---|---|---|---|---|
| California Sheephead | *Yes* | *Yes* | *no* | *no* | *yes* |
| Sea bass | *No* | *No* | *yes* | *no* | *yes* |
| Whitefish | *No* | *No* | *yes* | *no* | *yes* |
| Lobster | *Yes* | *No* | *yes* | *no* | *yes* |
| Abalone | *Yes* | *Yes* | *yes* | *yes* | *yes* |
| Sea cucumber | *No* | *No* | *yes* | *yes* | *yes* |
| Sea urchin | *Yes* | *Yes* | *yes* | *yes* | *yes* |
| Giant kelp | *Yes* | *Yes* | *no* | *no* | *no* |

**Supplementary Table 2.** Data sources and methods used to map intertidal, estuary, and subtidal marine habitats for the four subregions in the Southern California Bight.

| Region | Source | Description |
|---|---|---|
| Southern California Bight | [32] | Used existing giant kelp forest persistence maps for the California Bight based on Landsat images for 1984-2018. There is no giant kelp forest at Guadalupe Island. |
| | [33] | Used existing submarine canyon polygons for the California Bight. There are no seamounts or guyots in the 12 nm in the region. |
| Southern California | California Department of Fish and Wildlife | Used existing maps for southern California intertidal, estuary, and subtidal marine habitats |
| | [34] | Used existing tidal flat maps for Southern California We used the maximum coverage detected by Landsat images for 2008-2016 |
| Northern Baja California | [35] | Used existing maps for northern Baja California intertidal, estuary, and subtidal marine habitats |
| | Mapped for this project | We digitalized the eelgrass for Punta Banda and Bahía San Quintín Estuaries using maximum likelihood classification in ENVI (version 5.5.2, L3Harris Geospatial, Melbourne, FL, USA) based on a training set developed from on the ground presence-absence surveys and using low-altitude drone imagery |
| | Mapped for this project | We digitalized eelgrass (*Zoostera marina*) for Punta Banda and Bahía San Quintín Estuaries using maximum likelihood classification in ENVI (version 5.5.2, L3Harris Geospatial, Melbourne, FL, USA) based on a training set developed from in situ presence-absence surveys and using low-altitude drone imagery |
| Central Baja California | Pronatura | We obtained salt marshes, tidal flats, channels, and eelgrass for central Baja California |
| | Mapped for this project using Google earth | Used to digitalize the coastline for continent and islands and to differentiate between sand beach and rocky intertidal habitats in central Baja California and Guadalupe Island region. We digitalized rocky subtidal (those areas that we could interpret as rocky reef, kelp forest (maximum surface cover from the available images), and islets and assumed that the rest was sandy subtidal habitat |
| Northern and Central Baja California and Guadalupe Island | [36] | We extracted the bathymetric depth contours of 30, 100, and 200 m to obtain the different depth ranges for the three regions in Baja California |

**Supplementary Table 3.** List of marine habitats for intertidal, estuary, subtidal, and deep-sea habitats, and their coverage in each sub region in the Southern California Bight.

| **Habitat category** | **Habitat type** | **Southern California** | **Northern Baja California** | **Central Baja California** | **Guadalupe Island** | **California Bight total** |
|---|---|---|---|---|---|---|
| Intertidal (km) | Coastal Beaches | 405.64 | 297.45 | 384.61 | - | 1087.7 |
| | Coastal Rock | 92.90 | 261.85 | 340.00 | - | 694.75 |
| | Island Beaches | 194.2 | 3.5 | 60.06 | 4.64 | 262.4 |
| | Island Rock | 312.34 | 58.26 | 199.95 | 126.21 | 696.76 |
| Subtidal Sand ($km^2$) | 0-30m | 2116.53 | 1590.50 | 2650.64 | 50.34 | 6378.01 |
| | 30-100m | 5414.90 | 3202.06 | 8311.86 | 60.04 | 16988.86 |
| | 100-200m | 3511.69 | 958.76 | 2555.11 | 66.73 | 7092.29 |
| | Over 200m | 25580.27 | 4132.03 | 2864.81 | 3350.73 | 36227.84 |
| Subtidal Rock ($km^2$) | Coastal 0-30m | 12.92 | 33.29 | 35.50 | - | 81.71 |
| | Coastal 30-100m | 22.1 | ND | ND | ND | 22.1 |
| | Coastal 100-200m | 1.48 | ND | ND | ND | 1.48 |
| | Over 200 m | 2.02 | ND | ND | ND | 2.02 |
| | Island_0-30m | 20.19 | 4.07 | 22.68 | 20.34 | 67.28 |
| | Island_30-100m | 88.86 | ND | ND | ND | 88.86 |
| | Island 100-200m | 14.28 | ND | ND | ND | 14.28 |
| | Island Over 200 m | 140.94 | ND | ND | ND | 140.94 |
| Persistent Kelp (*Macrocystis pyrifera*) ($km^2$) | Coastal_Low | 43.77 | 54.91 | 8.95 | - | 107.63 |
| | Coastal_Mid | 29.74 | 35.20 | 10.14 | - | 75.08 |
| | Coastal_High | 9.11 | 12.58 | 10.85 | - | 32.54 |
| | Island_Low | 70.11 | 2.34 | 15.38 | - | 87.83 |
| | Island_Mid | 59.45 | 2.98 | 20.33 | - | 82.76 |
| | Island_High | 18.59 | 0.76 | 11.21 | - | 30.56 |
| Seagrass ($km^2$) | Surfgrass (*Phyllospadix sp.*) | 166.98 | ND | ND | ND | 166.98 |
| | Island Surfgrass (*Phyllospadix* sp.) | 188.50 | ND | ND | ND | 188.50 |
| | Eelgrass (*Zostera marina*) | 20.50 | 13.63 | 146.2 | - | 180.33 |
| Estuary ($km^2$) | Channel | 66.50 | 14.40 | 727.03 | - | 807.93 |
| | Tidal flats* | 4.77 | 19.12 | 106.83 | - | 130.72 |
| | Salt marshes | 15.65 | 18.61 | 156.47 | - | 190.73 |
| Others ($km^2$) | Submarine Canyons | 2872.65 | 1218.32 | 708.69 | 33.07 | 4832.73 |
| | Coastal Islet | 0.05 | 0.16 | 0.30 | - | 0.51 |
| | Island Islet | 0.73 | 0.04 | 0.39 | 0.15 | 1.28 |

**Supplementary Table 4.** Planktonic larval duration (PLD) and spawning periods obtained from the scientific literature and used during the oceanographic modelling of larval dispersal for four focal species. See main text for explanation about calculating the reduction of PLD under future climate change scenario. Seasons correspond to the Northern hemisphere. We also show the graph density (D), average link weight (AW) and average local retention (LR) for each of the larval dispersal networks. Note that for some focal species we used PLD information based on more than one species.

| **Focal species** | **Focal species used** | **PLD (days)** | **Spawning period** | **PLD Current scenario** | **PLD Future scenario** | **References** |
|---|---|---|---|---|---|---|
| Sea urchin | *M. franciscanus*<br>*Strongylocentrotus purpuratus* | 40-90 | Winter | 56 days<br>D: 17.99%<br>AW: 1.356<br>LR: 6.09% | 42 days<br>D: 10.58%<br>AW: 0.074<br>LR: 6.77% | [37-40] |
| Sheephead | *S. pulcher* | 39 | Fall | 42 days<br>D: 11.77%<br>AW: 2.214<br>LR: 6.80% | 28 days<br>D: 5.31%<br>AW: 0.205<br>LR: 7.17% | [41,42] |
| Abalone | *H. corrugata*<br>*H. fulgens* | 5-10 | Fall, Winter, Spring | 7 days<br>D: 4.26%<br>AW: 9.03<br>LR: 24.87% | 5 days<br>D: 1.57%<br>AW: 0.648<br>LR: 36.64% | [43,44] |
| Giant kelp | *Macrocystis pyrifera* | hours to 2 days | Winter, Spring, Summer, Fall | 2 days<br>D: 2.16%<br>AW: 14.753<br>LR: 45.33% | 2 days<br>D: 1.64%<br>AW: 0.589<br>LR: 37.22% | [45] |

**Supplementary Table 5.** Simulation parameters for the six focal species used in the deterministic, discrete-time logistic growth model with spatially implicit reserve and fishing zones. Note that for some focal species we used information of catch based on more than one species caught in the region.

| Focal species | Focal Species used | r | Fmsy |
|---|---|---|---|
| Whitefish | *Caulolatilus princeps* | 0.240 | 0.081 |
| Sea bass | *Paralabrax nebulifer* | 0.281 | 0.081 |
| Lobster | *P. interruptus* | 0.119 | 0.079 |
| Abalone | *H. assimilis, H. corrugata, H. cracherodii, H. fulgens, H. rufescens, H. sorenseni, H. sorenseni* | 0.079 | 0.073 |
| Sea urchins | *M. franciscanus, S. purpuratus* | 0.245 | 0.081 |
| Sea cucumber | *Parastichopus parvimensis* | 0.687 | 0.281 |

**Supplementary Table 6.** Species used to run the stochastic version of discrete-time logistic growth which includes the probability of marine heatwaves and the effect on biomass. Note that for some focal species we used density information of monitoring data[46] based on more than one species.

| Focal species | Focal species used |
|---|---|
| Abalone | *H. assimilis, H. corrugata, H. fulgens, H. rufescens, H. sorenseni* |
| Sea uchins | *M. franciscanus, S. purpuratus* |
| Sea cucumbers | *P. californicus, P. parvimensis* |

**Supplementary Table 7.** Catch in tonnes from 2000-2018 for six small scale fishery species, except sea cucumbers (2000-2015) for northern Baja California, central Baja California, and Guadalupe Island.

| Focal species | Focal Species used | Northern Baja California | Central Baja California | Guadalupe Island |
|---|---|---|---|---|
| *Sheephead* | *S. pulcher* | 1,350 | 207 | 0 |
| *Sea bass* | *Paralabrax clathratus* | 827 | 1520 | 0 |
| *Lobster* | *P. interruptus* | 1,993 | 28,787 | 131 |
| *Abalone* | *H.* spp. | 6.6 | 7,872 | 598 |
| *Sea cucumbers* | *Apostichopus and Parastichopus* spp. | 2,058 | 782 | 7 |
| *Sea urchin* | *M. franciscanus* | 35,509 | 3,091 | 0 |

**Supplementary Table 8.** Minimum area or linear coastal length required for a habitat to be considered a replicate inside a marine reserve, based on the Marine Life Protection Act Initiative[47].

| Habitat | Minimum threshold for replication |
|---|---|
| Rocky intertidal | 0.97 km |
| Sandy beach | 1.77 km |
| Giant Kelp | 1.77 km |
| Subtidal rock (0-30 m) | 1.77 km |
| Subtidal rock (30-100 m) | 0.52 $km^2$ |
| Subtidal rock (100-3000 m) | 0.52 $km^2$ |
| Subtidal sand (0-30 m) | 1.77 km |
| Subtidal sand (30-100 m) | 12.9 $km^2$ |
| Subtidal sand (100-3000 m) | 18.1 $km^2$ |
| Coastal marsh | 0.10 $km^2$ |
| Eelgrass | 0.10 $km^2$ |
| Estuary | 0.31 $km^2$ |

**Supplementary Table 9**. Spatial and temporal scales of movements between juvenile and adult habitats, and method used to determine movements.

| Common name | Scientific name | Spatial scale (km) | | | | Temporal scale | Juvenil habitat | Adult habitat | Habitat use | Method | Source |
|---|---|---|---|---|---|---|---|---|---|---|---|
| | | $10^0$ | $10^2$ | $10^3$ | $10^4$ | | | | | | |
| **Sharks** | | | | | | | | | | | |
| White shark | *Carcharodon carcharias* | - | - | - | - | >3 months | Point Conception, CA.to BCS coast (Sebastian Vizcaino Bay, BC) | Central California islands and Guadalupe Island | Nursery, foraging | 3 | [48]*; [49]*; [50]*, [51]* |
| Salmon shark | *Lamna ditropis* | - | - | - | | >1 year | | Open ocean off BC | Breeding and foraging | 3 | [52]* |
| Blue shark | *Prionace glauca* | - | - | - | | 1 month | | Along the BC and BCS coast | Foraging and feeding | 3, 4 | [53]*; [54]* |
| Basking shark | *Cetorhinus maximus* | - | - | - | - | >3 months | | Point Conception, CA. to north of BC and Hawaii | Foraging | 3, 4 | [55]*; gtopp.org* |
| Shortfin mako shark | *Isurus oxyrinchus* | - | - | - | - | 1 year | | Along the BC and BCS coast | Foraging and feeding | 3, 4 | [56]*; [54]* |
| Thresher shark | *Alopias vulpinus* | - | - | - | | >6 months | Punta Eugenia, BC. to Morro Bay, CA | - | Foraging | 4 | [57]* |
| Smooth hammerhead shark | *Sphyrna zygaena* | | | | | | | | | | gtopp.org* |
| **Bony fishes** | | | | | | | | | | | |
| Swordfish | *Xiphias gladius* | | | | | | | | | | gtopp.org* |
| White seabass | *Atractoscion nobilis* | - | - | | | >2 years | | Half Moon Bay, CA and La Salina, BC | Feeding and spawning | 2 | [58] |
| Yellowtail | *Seriola lalandi* | - | - | | | >6 months | Offshore | Inshore of Santa Barbara to the BC coast | Foraging and feeding | 2 | [59]* |
| Bluefin tuna | *Thunnus orientalis* | - | - | - | | >3 months | California−Oregon border and to Magdalena Bay, BCS | California−Oregon border and to BC-BCS border | Feeding | 3, 4 | [60]*; [61]* |
| Yellowfin tuna | *Thunnus albacares* | - | - | - | | >1.5 years | | Along BCS coast | Foraging and feeding | 4 | [62] |
| California scorpionfish | *Scorpaena guttata* | - | - | | | >1.5 years | | Mexico-USA border | Spawning | 2 | [63] |
| Vermilion rockfish | *Sebastes miniatus* | - | - | | | >1.5 years | | Mexico-USA border | Spawning | 2 | [63] |
| **Sea Birds** | | | | | | | | | | | |

| | | | | | | | | | | | |
|---|---|---|---|---|---|---|---|---|---|---|---|
| Laysan albatross | *Phoebastria immutabilis* | | | | | | | Guadalupe Island | Breeding | 3 | [64]*; [65]* |
| Black-footed albatross | *Phoebastria nigripes* | - | - | - | | >2 months | | Offshore San Quintin, BC | Post-breeding dispersal | 3 | [66] |
| Pink-footed shearwater | *Ardenna creatopus* | - | - | - | - | >6 months | | Along the BC and BCS coast | Non- breeding season | 3 | [67]* |
| Sooty shearwater | *Puffinus griseus* | - | - | - | - | | | | | 3 | [68]* |
| Black-vented Shearwater | *Puffinus opisthomelas* | - | - | | | >15 days | | El Vizcaino Bay to the southern Gulf of Ulloa | Breeding, foraging and resting | 3,4 | [69]* |
| **Marine mammals** | | | | | | | | | | | |
| Blue whale | *Balaenoptera musculus* | - | - | - | - | >6 months | | Along the BC and BCS coast | Feeding, breeding and foraging | 3 | [70]*; [71]* |
| Grey whale | *Eschrichtius robustus* | - | - | - | - | | | | | | [72]* |
| Northern elephant seal | *Mirounga angustirostris* | - | - | - | | | | | | 3, 4 | [73]*, gtopp.org* |
| California sea lion | *Zalophus californianus* | | | | | | | | | | gtopp.org* |
| Bottlenose dolphin | *Tursiops truncatus* | - | - | | | >15 days | | Punta Colonet to Point Conception | Feeding | 1 | [74] |
| **Sea Turtles** | | | | | | | | | | | |
| Leatherback turtle | *Dermochelys coriacea* | - | - | | | >1 year | | From Washinton to Ensenada | Foraging | 3 | [75]* |
| Loggerhead sea turtles | *Caretta caretta* | - | - | - | | >1 year | | Along the BC and BCS coast | Foraging | 3 | [76] |

*Sources used for Figure 4.4.

**Method: (1) Photoidentification or direct counting, (2) conventional tags or spaghetti, (3) satellite tags, and (4) archival tags

**Supplementary Table 10.** Results from the linear models comparing scaled yearly density change for years with marine heatwaves and years without for abalone, for sea cucumber, and for sea urchin.

| Factor | DF | Sum of Sq | Mean Sq | F value | Pr(>F) |
|---|---|---|---|---|---|
| **Abalone** | | | | | |
| Threshold | 1 | 6.13 | 6.1303 | 4.5347 | 0.035 |
| **Sea cucumber** | | | | | |
| Threshold | 1 | 9.531 | 9.5308 | 6.5049 | 0.012 |
| **Sea urchin** | | | | | |
| Threshold | 1 | 12.96 | 12.957 | 8.4855 | 0.004 |

#### Supplementary Figures

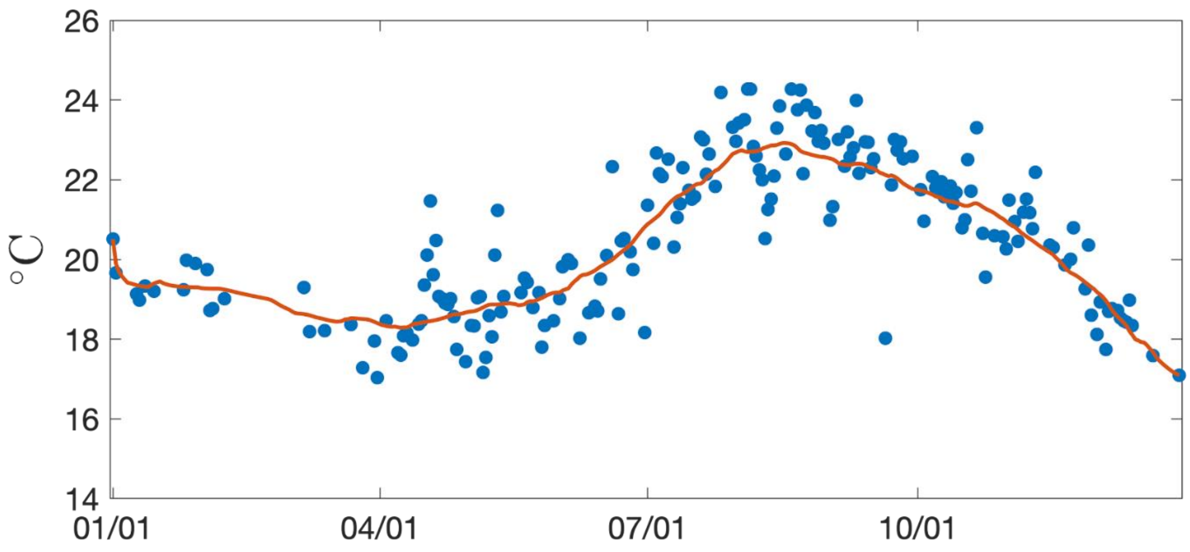


**Supplementary Fig 1.** Example of SST derived at Punta Prieta, Isla Natividad in central Baja California for 2001. The red line represents the annual climatology for Punta Prieta, and the blue points the daily SST. SST variability was derived by computing the variance relative to the moving mean of the annual cycle.

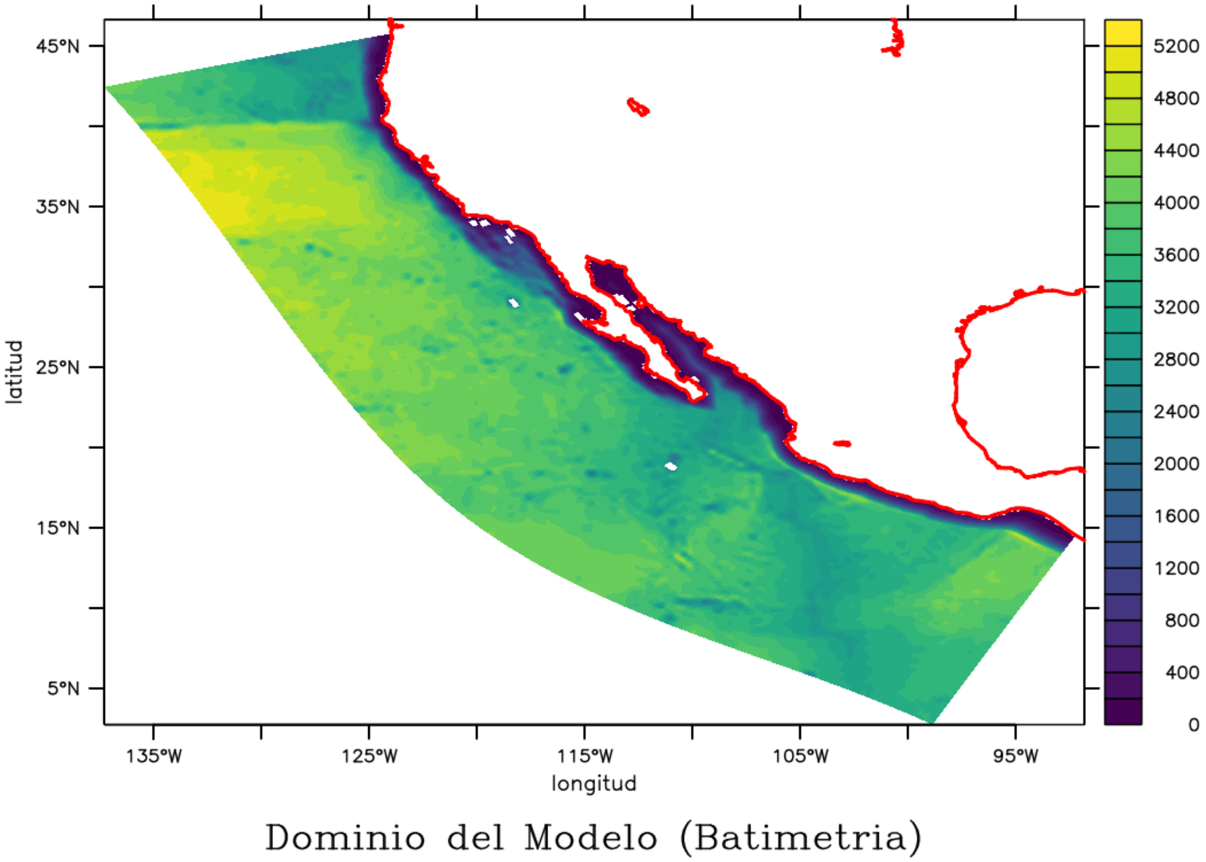


**Supplementary Fig 2.** Spatial domain of the ROMS model used to study the larval connectivity between California USA and Mexico.

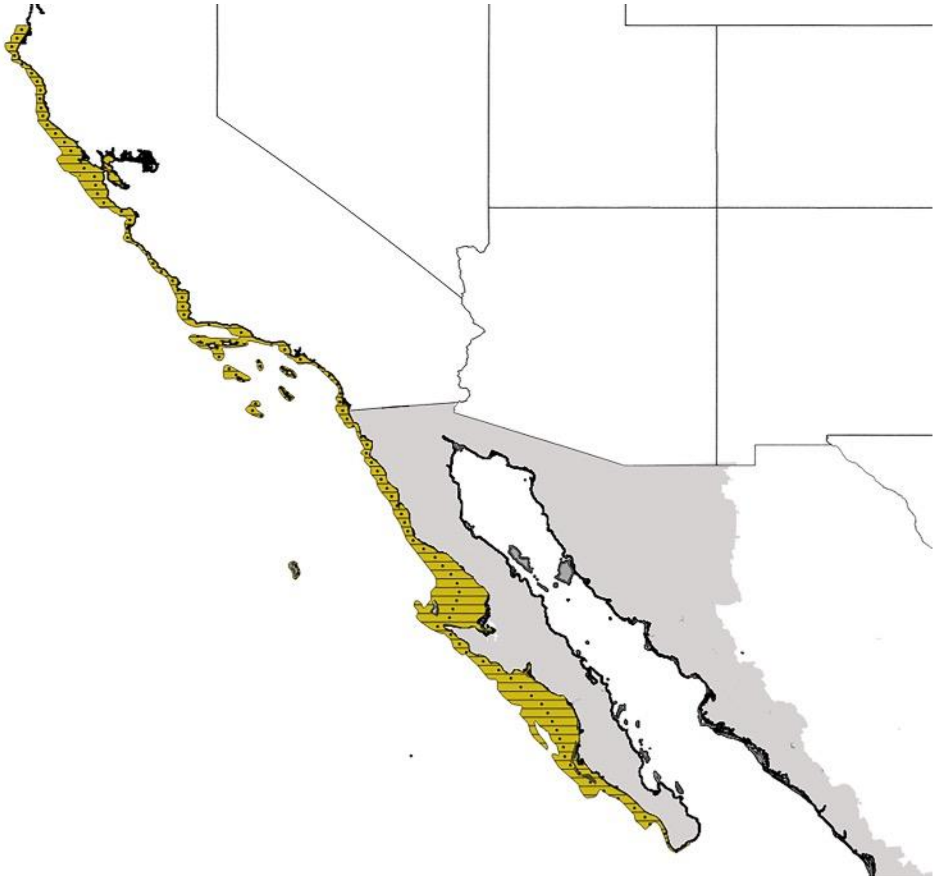

**Supplementary Fig 3.** Spatial units used in the analyses of larval connectivity between California USA and the Baja California Peninsula. The 115 units are defined by the coast and the 200 m isobath.

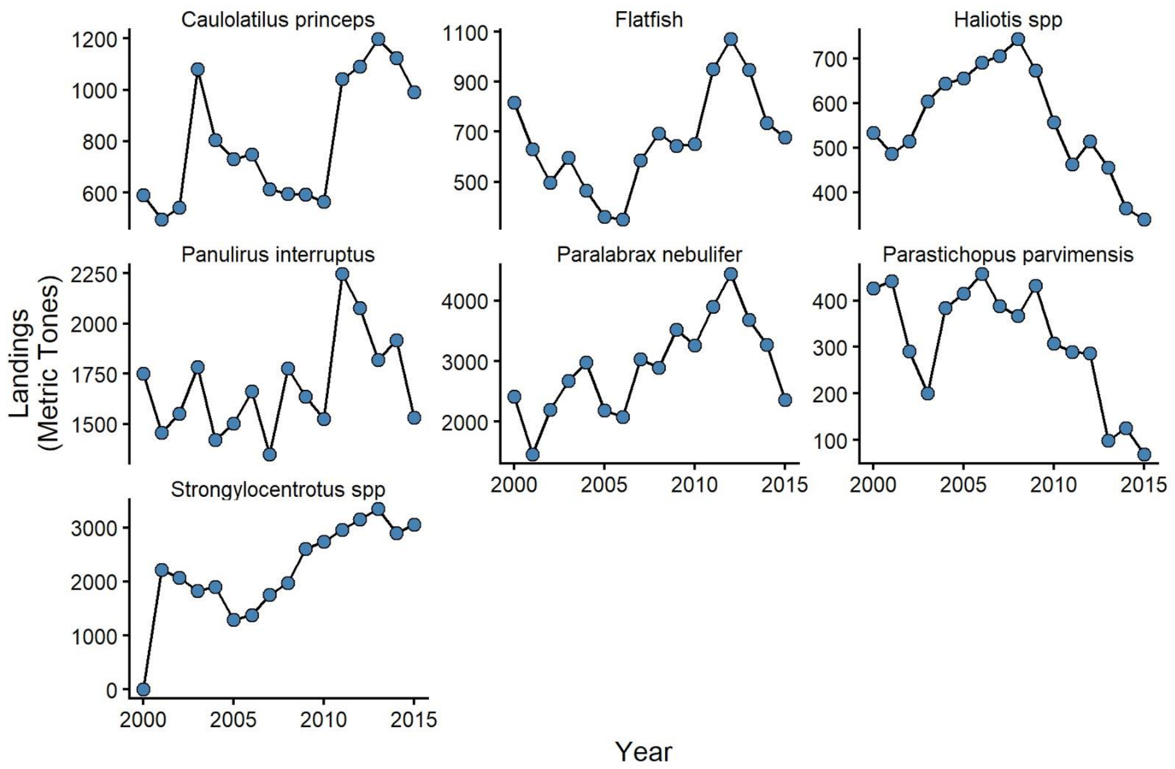


**Supplementary Fig 4.** Timeseries of landings (tons) for each species used in the deterministic, discrete-time logistic growth model with spatially implicit reserve and fishing zones. Data come from CONAPESCA landing receipts (2000 - 2015).

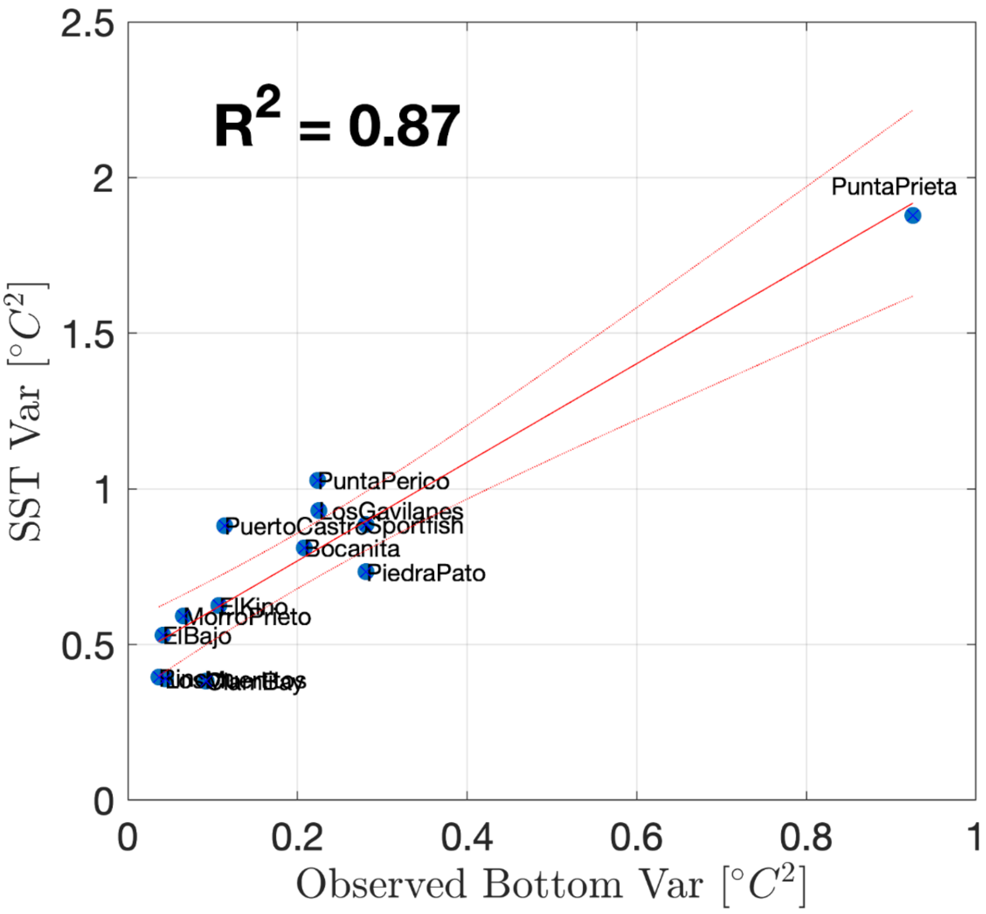


**Supplementary Fig 5.** SST temperature variability compared to observed bottom temperature variance at sub-daily periods.

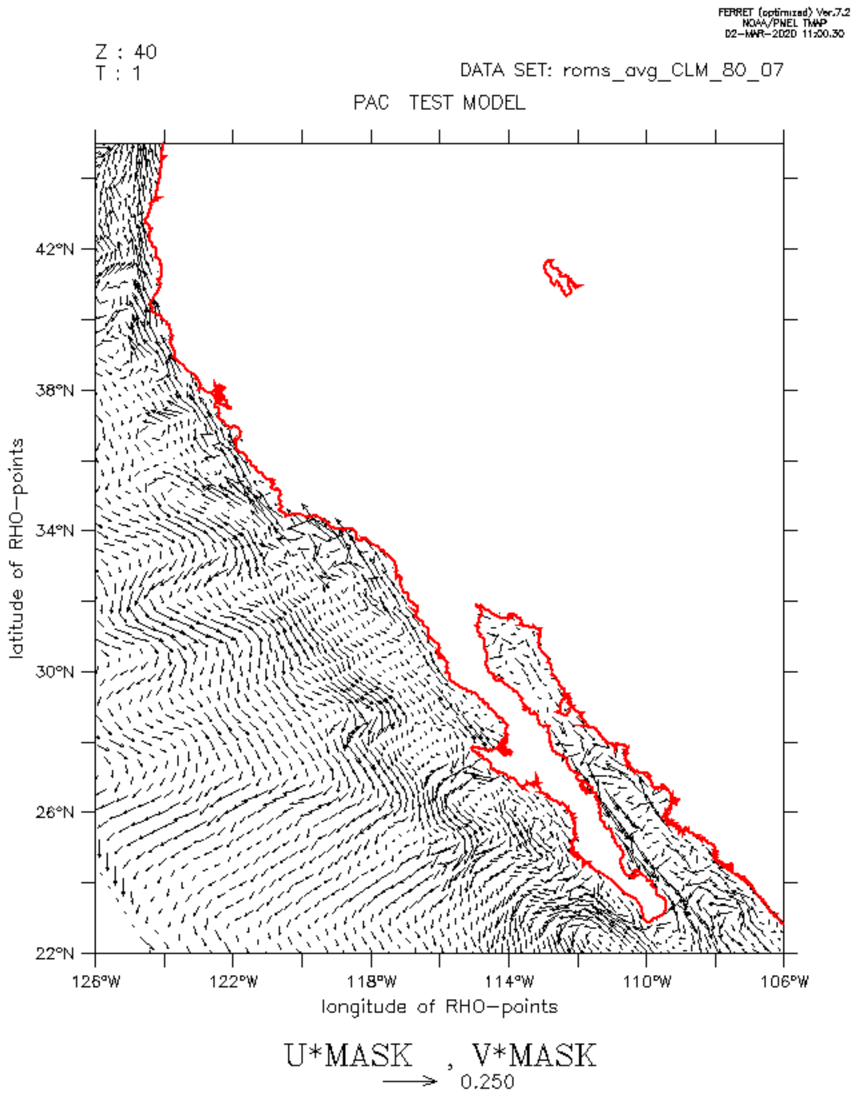


**Supplementary Movie 1.** Animation showing vectors of current speed and direction in the study area between California (USA) and the Baja California Peninsula (Mexico).
